# iLipidome: enhancing statistical power and interpretability using hidden biosynthetic interdependencies in the lipidome

**DOI:** 10.1101/2024.05.16.594607

**Authors:** Wen-Jen Lin, Austin W.T. Chiang, Evanston H. Zhou, Chenguang Liang, Chia-Hsin Liu, Wen-Lung Ma, Wei-Chung Cheng, Nathan E. Lewis

## Abstract

Numerous biological processes and diseases are influenced by lipid composition. Advances in lipidomics are elucidating their roles, but analyzing and interpreting lipidomics data at the systems level remain challenging. To address this, we present iLipidome, a method for analyzing lipidomics data in the context of the lipid biosynthetic network, thus accounting for the interdependence of measured lipids. iLipidome enhances statistical power, enables reliable clustering and lipid enrichment analysis, and links lipidomic changes to their genetic origins. We applied iLipidome to investigate mechanisms driving changes in cellular lipidomes following supplementation of docosahexaenoic acid (DHA) and successfully identified the genetic causes of alterations. We further demonstrated how iLipidome can disclose enzyme-substrate specificity and pinpoint prospective glioblastoma therapeutic targets. Finally, iLipidome enabled us to explore underlying mechanisms of cardiovascular disease and could guide the discovery of early lipid biomarkers. Thus, iLipidome can assist researchers studying the essence of lipidomic data and advance the field of lipid biology.

## Introduction

Lipids are an essential energy reservoir, important modulators of signaling, and fundamental building blocks of cellular membranes. Advances in mass spectrometry (MS) can link lipids to phenotypes^1–4^. Lipidomics analyses often focus on a handful of lipids showing the largest changes in abundance for biological insight^5–7^. However, there are several limitations to this approach. First, after correcting for multiple hypothesis testing, individual lipids might not meet statistical significance due to the biological variability across samples or the noise generated by lipidomic technologies^8^. Second, in other cases, one may obtain a long list of statistically significant lipids without a clear unifying biological theme; thus *ad hoc* interpretation is required. Third, focusing on individual lipids may miss crucial concerted changes shared across whole pathways. Fourth, since most conventional lipidomics analyses overlook the interconnections among measured lipids, it remains challenging to identify the genetic source of lipidomic changes. For these difficulties, analysis of the data in the context of the lipid molecular network could be a promising solution.

Essentially, a lipid molecular network encoding biosynthetic information could provide a comprehensive understanding of the complex metabolic processes and enable us to explore the perturbed pathways and reactions behind lipid profiles. Furthermore, lipids sharing biosynthetic pathways usually participate in similar biological processes as unique structural subgroups. For instance, it was found that persistent EGFR signaling increased saturated phosphatidylcholine (PC) while concomitantly reducing saturated lysophosphatidylcholine (LPC) in glioblastoma (GBM) cells^2^. Indeed, lysophosphatidylcholine acyltransferase 1 (LPCAT1), which converts LPC to PC, is critical to lipid remodeling and supports oncogenic signaling. Another study employed lipidomics to reveal striking changes to the linoleic acid (LA) to arachidonic acid (AA) metabolism axis upon coronavirus infection^9^. Thus, modulation of the LA-AA axis could help combat emerging coronaviruses through exogenous supplement of LA or AA or inhibition of cytosolic phospholipase A2α (cPLA2α). These studies highlight the value of a lipid network accounting for the interconnections and interdependence of lipidomics data.

While many lipidomics analysis tools exist, most do not account for the pathway context nor interdependencies of measured lipids. Web servers such as LipidSig^5^ and LipidSuite^6^ allow researchers to interrogate the associations between diverse lipid characteristics and biological phenotypes. Yet, analysis of single lipids is usually limited by low statistical power after false discovery rate (FDR) correction. Moreover, while lipid characteristics encode structural similarity among lipids, they do not account for their interconnections in the lipid biosynthetic pathways. BioPAN introduces a lipid network analysis approach^10,11^ and enables interpretation of altered pathways. Nevertheless, this method cannot address the interdependence in raw lipidomics data, making the following computation influenced by highly variable or sparsely measured lipids and therefore impacting the accuracy of analyses. Thus, the interconnections and non-independence in lipidomics data require a new approach to more precisely analyze the altered pathways and the genetic source of a lipid profile.

To address these challenges, we present iLipidome, a novel systems biology-based approach to analyze lipidomics data. Importantly, iLipidome accounts for biosynthetic steps and interdependences of experimentally measured lipids. It decomposes each lipid into intermediate substructures that can aggregate slight changes in lipid profiles based on their shared biosynthetic network^12^. This method significantly improves statistical power, reduces data sparsity, and increases biosynthetic network coverage in lipidomics data analysis. We also show its value in hierarchical clustering and lipid enrichment analysis. Furthermore, the substructure network encoding biosynthetic information can explore the mechanisms behind changes in lipid abundance. Here, we pinpointed the genetic sources of lipidomic changes following docosahexaenoic acid (DHA) supplementation. The hidden lipid interactions revealed by iLipidome also enabled us to identify the substrate preferences of LPCAT1 and the therapeutic targets for GBM. Lastly, we demonstrated such analyses could reveal the potential mechanisms and guide the early biomarker development for cardiovascular disease (CVD). Collectively, iLipidome facilitates systems-level comparison of lipid profiles with lipid biosynthetic information efficiently and provides a deeper insight into complex lipidomic alterations across samples.

## Result

### High variability in lipid measurements introduces challenges in conventional lipidomics analysis

Current lipidomics analysis strategies mainly focus on individual lipids or lipid characteristics. However, since many lipids share biosynthetic steps and enzymes, changes in one lipid may impact the abundance of many others. To understand this dependency and how it impacts lipidomics analysis, we first applied conventional lipidomics analysis methods on a dataset studying the effect of DHA (22:6 omega-3) treatment on rat basophilic leukemia (RBL) cells^3^.

The general differential abundance analysis showed the number of significant lipid species drastically decreases after correcting for multiple hypothesis testing (**Fig. 1a**). This stems from the high variability and sparsity^8^ in detected lipids across samples, wherein few lipids (90/444=20.3%) were measured in all samples (**Fig. 1b**). Many studies have confirmed that the *de novo* synthesis of unsaturated fatty acids (FAs) in cells will be inhibited by DHA supplementation^3,13,14^. Nonetheless, the high variability impeded the discovery of this important FA remodeling signature in the enrichment analysis after FDR correction (**Fig. 1c**). Practically, one can categorize lipids into different subgroups in terms of specific lipid characteristics (e.g., FA chain length and double bond number) to improve the statistical power^5,6^. Thus, we employed a similar analysis and found the expected rise in the categories of chain length 22 and double bond 6 (p-value < 0.001), corresponding to the feature of DHA (**Fig. 1d,e**). Further, the chain length-double bond combined analysis unveiled a more precise pattern of FA change (e.g., increase in 16:0 and 22:6 FAs (p-value < 0.01) or decrease in 18:1 and 20:2 FAs (p-value < 0.05)) (**Fig. 1f**). These methods provide insights into shared lipid features and increase statistical power by aggregating lipids with similar structures. However, the aggregation does not consider the sequential connections of the lipids or FAs determined by biosynthetic pathways. It’s also difficult to identify genetic source of change, despite noticing a consistent decrease in abundance of a subgroup of unsaturated FAs (18:1, 20:1, and 20:2 FAs) (**Fig. 1f**). Thus, the conventional methods fail to balance the statistical power and the integrity of individual lipids at the same time, and they further pose a challenge in identifying the genetic basis of a global reprogramming of the lipidome. To address these challenges, we developed iLipidome, a substructure-based approach that uses the lipid biosynthetic pathways for data analysis to solve the non-independence issue in lipidomics data, provide interpretable biosynthetic information, and explore impacted pathways and enzyme catalyzed reactions.

**Fig 1:**
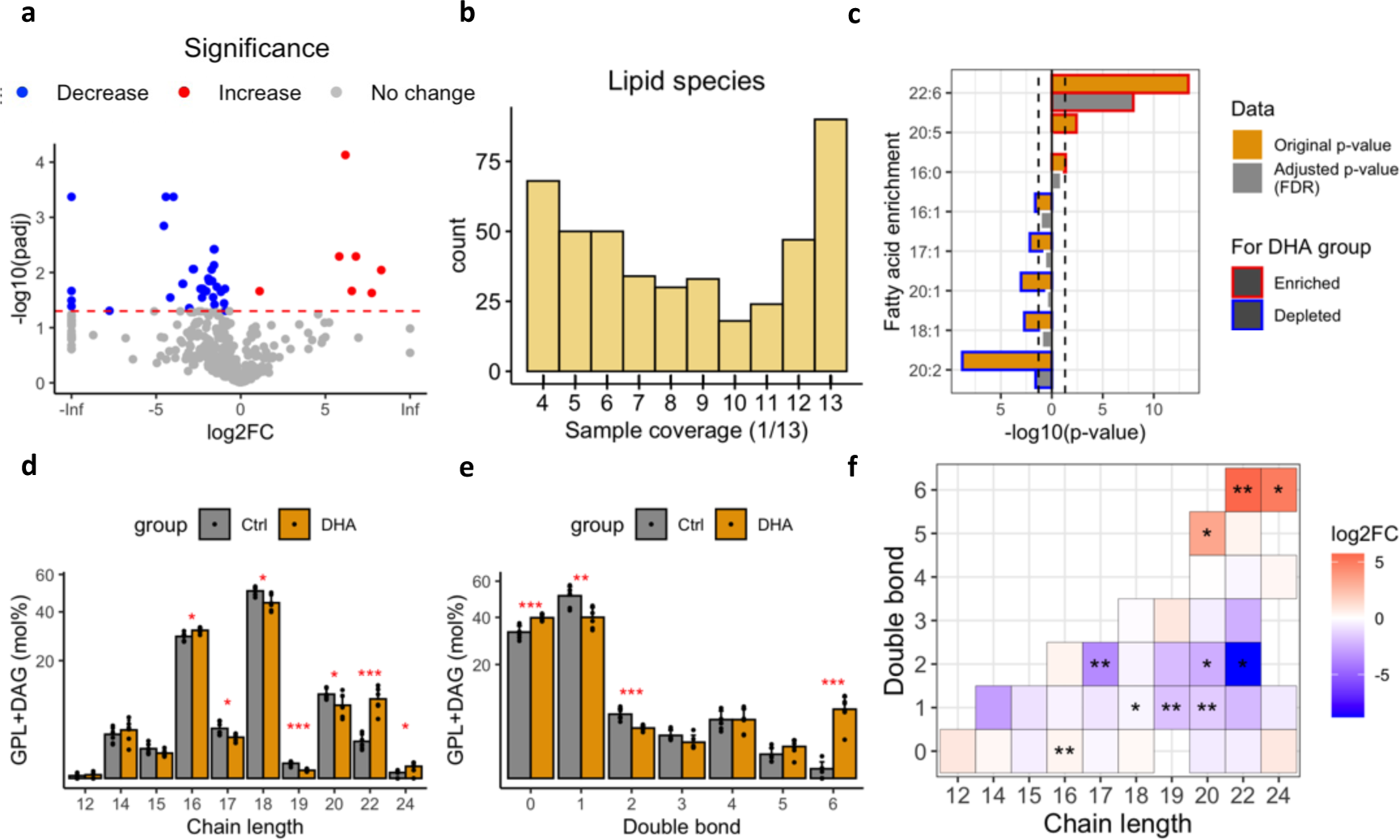
Limitations in conventional lipidomics analysis methods. **a**, The volcano plots show the differential abundance of lipid species based on adjusted p-values in the comparisons between 7 controls and 6 DHA-treated RBL cell samples. **b**, The sample coverage of lipid species is quantified as the frequency of a lipid species detected in 13 samples. Many lipid species show low overlap in the raw lipidomics data. The sparsity and non-independence in lipid profiles are not corrected in differential expression analysis, which contributes to the low statistical power. **c**, FA enrichment analysis for the statistically significant lipid species before and after false discovery rate (FDR) correction. The significantly enriched and depleted FAs were outlined in red and blue, respectively. **d-f**, We calculated (**d**) FA chain length profile, (**e**) lipid unsaturation (double bond) profile, and (**f**) the two-characteristics combined heatmap in membrane glycerophospholipids (GPLs) and diacylglycerols (DAGs) in the DHA dataset. DHA supplementation not only leads to a dramatic increase in its corresponding 22:6 FA but also causes obvious remodeling in the other FAs. However, these methods cannot explain the interconnected nature of altered lipids and the genetic source of changes. Two-tailed Student’s t-tests were used to calculate the p-values in (**a**) and (**d-f**). Benjamini–Hochberg p-value correction was used for all analyses. The dashed lines in (**a**) and (**c**) indicate p-value=0.05. *p<0.05, **p<0.01, ***p<0.001.

### iLipidome decomposes lipid profiles into biosynthetic precursors to account for interdependencies during analyses

iLipidome is a systems biology-based method to analyze lipidomics data by decomposing lipids into substructures, reconstructing biosynthetic network, and identifying critical pathways and genes/enzymes. First, we manually curated the FA and lipid class pathways from KEGG^15^, Reactome^16^, and recent lipid studies and reviews (**Fig. 2a**)^17–22^. Further, the iLipidome algorithm can combine the resulting lipid biosynthetic network with the LIPID MAPS database^23^ and the FA information encoded in lipids to decompose the lipids into three types of substructures: lipid class substructures, FA substructures or lipid species substructures, based on their biosynthetic routes (**Fig. 2b**). Specifically, the abundance of each measured lipid can be backpropagated to their biosynthetic intermediates, and summed when more than one lipid shares a common substructure. iLipidome, therefore, accounts for interdependencies and interconnections between lipids by searching for substructures (biosynthetic intermediates) of the experimentally measured lipids in lipidomics data. A FA or lipid can be synthesized from the *de novo* (endogenous) pathway using a basic structural unit, such as acetyl-CoA or glycerol-3-phosphate (G3P), or from an exogenous intermediate absorbed by a cell^24^. Additionally, changes in abundance of the measured lipids could emerge from increased/decreased uptake of related precursors or alterations of enzyme activities. Theoretically, absorbed lipid intermediates or activated enzymes will increase the products in their downstream pathways, while inhibiting upstream pathways. We therefore use the fold change of each substructure between two experimental groups to identify the potential source of altered lipids and further extract the substructures in one biosynthetic path (**Fig. 2c**). Following the rules above, each lipid in a lipidomics dataset can be transformed to their substructures, all of which will be further used to produce a substructure matrix recording the frequency of occurrence of each substructure (**Fig. 2d**). The substructure abundance can then be calculated using the substructure matrix and the lipid profiles. The summation of each substructure is not simply the amount of an intermediate but reflects the activity of biosynthetic steps required to form the current lipid profiles. Therefore, the non-independence challenge is addressed through the analysis of lipid substructures. Moreover, since the substructures from biosynthetic pathways are interconnected and can aggregate lipid signals, there is less concern of data sparsity (i.e., unmeasured lipid intermediates) to reconstruct the network from different lipid profiles (**Fig. 2e**). We further introduce a scoring system to identify essential pathways and reactions that are significantly activated or suppressed across conditions (**Fig. 2f**)^25^. iLipidome is presented as an R package and R-Shiny web-app (http://www.bioinfomics.org/iLipidome/) that allows for user-friendly and interactive data analysis. More detailed information on the reference pathway collection, substructure decomposition and extraction, and pathway and reaction scoring methods can be found in Materials and Methods.

**Fig 2:**
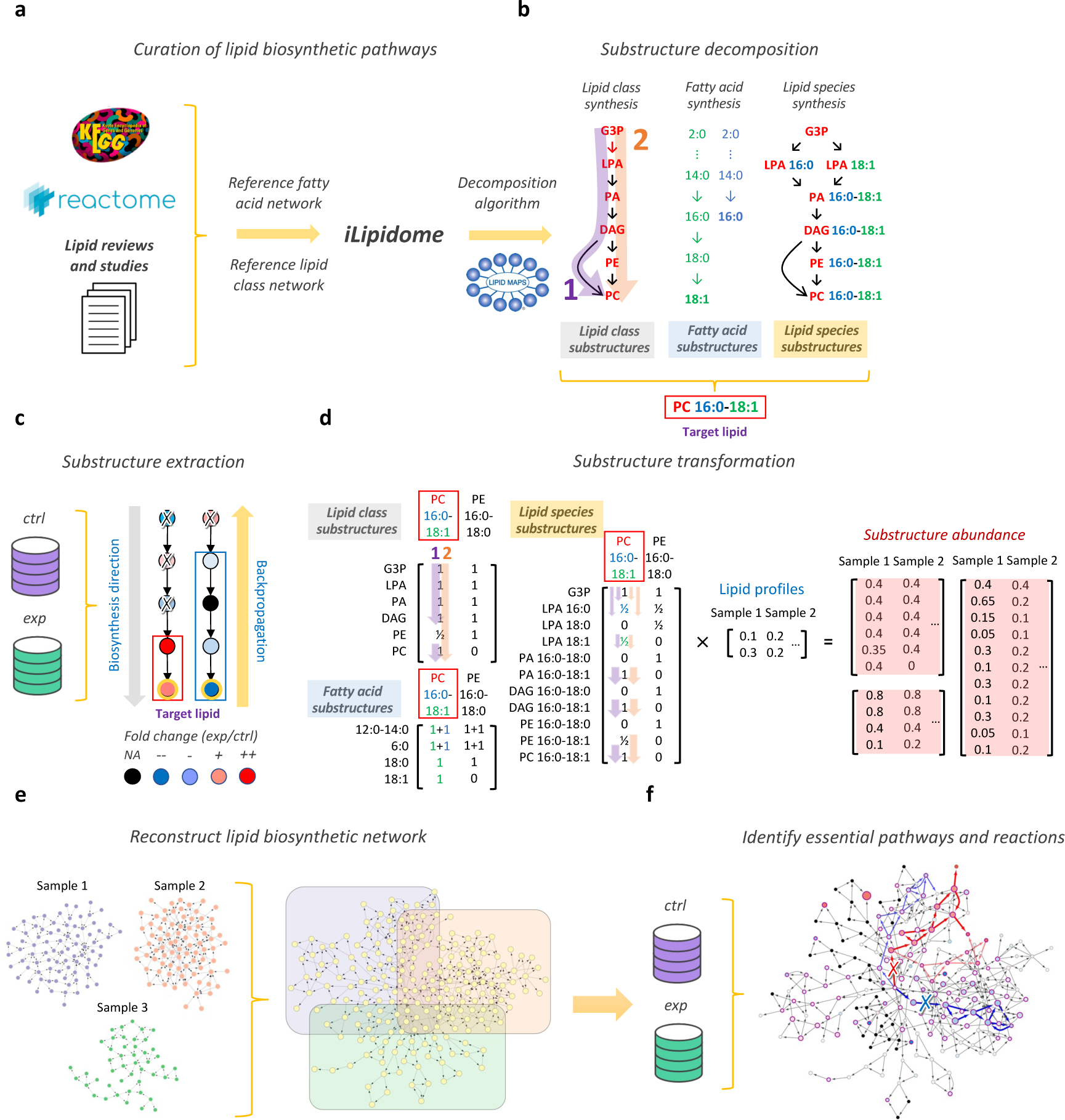
Schematic overview of the analysis workflow for iLipidome. **a**, The reference FA and lipid class biosynthetic pathways were manually curated from KEGG^15^, Reactome^16^, and recent lipid studies and reviews^17–22^. **b**, Lipids are decomposed to substructures (lipid class substructures, FA substructures or lipid species substructures) based on the reference biosynthetic pathways and the LIPID MAPS database^23^. **c**, To identify the potential source of altered lipids, substructure units in each biosynthetic route are further reduced through backpropagation and use of the fold changes between control and experimental groups (see Methods). For example, two lipid species (yellow nodes) were decomposed into their substructures. In the left pathway, the propagation stops at the third node due to the opposite fold change with the lipid species. Thus, only two substructures are kept (red rectangle). Likewise, four substructures are retained in the right pathway (blue rectangle). **d**, The presence frequency of each substructure for each lipid is recorded in a substructure matrix, which is further weighted by the lipid profiles and summed into the final substructure abundance matrix. **e**, The substructure profiles produced by each sample retain biosynthetic information, and these are combined to reconstruct a unified lipid biosynthetic network. **f**, iLipidome enables one to identify the essential pathways and reactions in the biosynthetic network. The significant representative pathways (color paths) and the highly perturbed reactions (crosses) are calculated and labeled in the network, with red and blue indicating activation and suppression, respectively. Edge colors and weights can indicate pathway importance.

### iLipidome increases the statistical power and reveals genetic source responsible for DHA-induced FA remodeling

We first applied iLipidome to the previous DHA dataset and compared the results from the original and substructure transformed data. By encoding biosynthetic context, iLipidome provides richer information (**Fig. 3a,b**) and significantly (p = 6.9e-04) enhanced statistical power than using raw FAs alone (**Fig. 3c**). Besides, it gives us the opportunity to examine altered pathways and enzymes based on lipidomics data only. In the FA substructure network, three significantly altered pathways can be clearly observed: (1) activation of omega-3 FA synthesis, (2) accumulation in saturated FAs, and (3) reduction in unsaturated FAs in the non-essential FA synthesis (**Fig. 3b,d**). Intriguingly, we identified several key desaturases and elongases, such as stearoyl-CoA desaturase 1 (SCD1), ELOVL1/2/3/5/7 (ELOVL Fatty Acid Elongase 1/2/3/5/7), and fatty acid desaturase 1/2 (FADS1/2), that may engage in the FA remodeling induced by DHA supplementation (**Fig. 3e**). Among them, SCD1 catalyzed reactions are decreased and found to locate exactly at the boundary between active and suppressed pathways, highlighting their central roles in overall non-essential FA remodeling (**Fig. 3b,e)**. In fact, down-regulation of SCD1 by polyunsaturated fatty acids (PUFAs) containing DHA has been validated in many studies^13,26^. It suppresses the crucial steps in FA unsaturation and counteracts DHA perturbations^3^. On the contrary, analysis of unprocessed lipidomics data using a conventional approach showed a lower statistical power, discovered only partially impacted pathways, and did not successfully identify critical SCD1 (**Fig. 1a,d and Extended Data Fig. 1a,b**).

**Fig 3:**
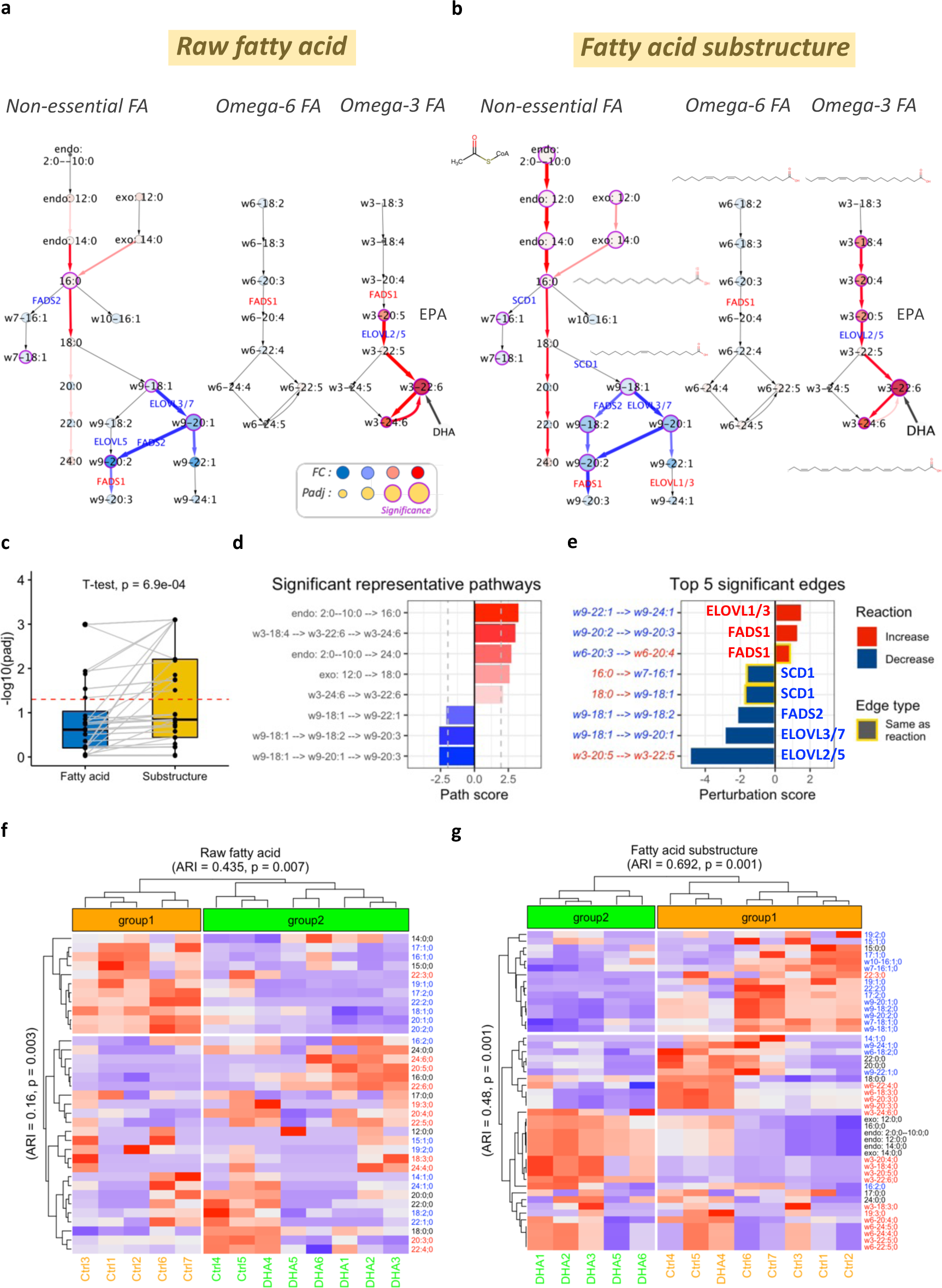
Analysis of FA substructures using iLipidome increases statistical power and elucidates FA remodeling and its genetic sources induced by DHA. **a,b**, The FA biosynthetic network derived from (**a**) the raw FA or (**b**) the iLipidome FA substructure data in RBL cells supplemented with DHA. Two-tailed Student’s t-tests with Benjamini–Hochberg correction were used to calculate the adjusted p-values. Nodes are filled according to log2 (fold change) and their sizes denote –log10 (adjusted p-value). If one node shows significant changes in abundance, its border will be marked as purple. The significant representative pathways and the highly perturbed edges are also labeled in the network, with red and blue indicating activation and suppression, respectively. Line width and color depth marks pathway importance. **c**, Boxplots reveal a significant improvement in the adjusted p-values of the substructures. **d**, Significantly activated and suppressed representative pathways in the network from panel (**b**). Pathways are simplified and represented by the first and the last metabolite. If different pathways have a same name, an additional metabolite is used to differentiate them. **e,** Top 5 significantly activated and suppressed edges (reactions) in the network from panel (**b**). Here, we removed reactions including DHA to avoid artifacts due to supplementation. One edge (reaction) is composed of two nodes (substrate and product), whose colors are defined as the fold changes of their abundance. Edge types including increase, decrease, and no change reflect the changes of the two associated nodes. Bars are outlined yellow if the edge type is same as the perturbation of reaction. **f**,**g**, Substructure profiles enable more precise clustering of biological samples and lipid characteristics. Heatmap of hierarchical clustering for (**f**) unprocessed FAs and (**g**) FA substructures in 13 samples. We divided samples into two groups and measured the clustering similarity with the original labels. FA variables on the rows are colored according to the double bond number. Black, blue, and red denote FAs without a double bond, with 1 and 2 double bonds, and with 3 or more double bonds, respectively. FA variables are also separated into two groups with black and red combined into one category and calculated the clustering similarity. EPA, eicosapentaenoic acid; DHA, docosahexaenoic acid; w3, omega-3; w6, omega-6; w7, omega-7; w9, omega-9; endo, endogenous; exo, exogenous; SCD1, stearoyl-CoA desaturase-1; FADS1/2, fatty acid desaturase 1/2; ELOVL1/2/3/5/7, ELOVL Fatty Acid Elongase 1/2/3/5/7; ARI, adjusted Rand index.

Another interesting finding is the increased activity of eicosapentaenoic acid (EPA, 20:5 omega-3) synthesis (**Fig. 3b)**, which iLipidome suggests can be attributed to the inhibition of ELOVL2/5 (**Fig. 3e**). This result is consistent with many other studies, which suggest that DHA supplementation may slow down EPA metabolism and lead to an accumulation of EPA synthesized from alpha-linolenic acid (ALA, 18:3 omega-3) (**Fig. 3b**)^17,27–29^. Indeed, DHA feeding lowers the expression of ELOVL2/5 in mice^28,29^. Many other elongase involved reactions including ELOVL1/3/7 were also detected changed by DHA treatment (**Fig. 3e**). Generally, they are responsible for long chain saturated and monounsaturated FA elongation^30,31^. However, how they are reversely modulated by PUFAs is poorly studied in comparison to ELOVL2/5^32^.

Since the substructures summarize the biosynthetic context and highlight the difference across conditions, we further tested if the hidden biosynthetic similarities between lipids could benefit sample clustering. Indeed, FA substructures displayed a higher adjusted Rand index value (0.435 to 0.692) and a more consistent clustering on the DHA samples (**Fig. 3f-g**). Further, in the row clustering, the highly unsaturated FAs (3 or more double bonds) and saturated FAs were clearly separated from the other unsaturated FAs (1 and 2 double bonds), consistent with the previous conclusion of DHA-induced FA remodeling (**Fig. 3b,g**). We also observed DHA4 has less DHA incorporation, which can explain why it is an outlier and closer to control group when clustering (**Extended Data Fig. 1c,d**). Taken together, iLipidome successfully increases the statistical power and reliability for distinguishing samples and FA features correctly. More importantly, it accurately predicts the pivotal pathways and reactions that are hardly visible using conventional methods, providing more comprehensive insights into overall FA remodeling. These advantages can also be observed when we decomposed lipids to species substructures (**Fig. 4 and Extended Data Fig. 2 and Supplementary Text 1**). The additional statistical power relies on improvement of data sparsity and enhance following enrichment analysis. Finally, iLipidome exhibited a higher network coverage and greatly increased the interpretability of lipidomics data by reconstructing its biosynthesis.

**Fig 4:**
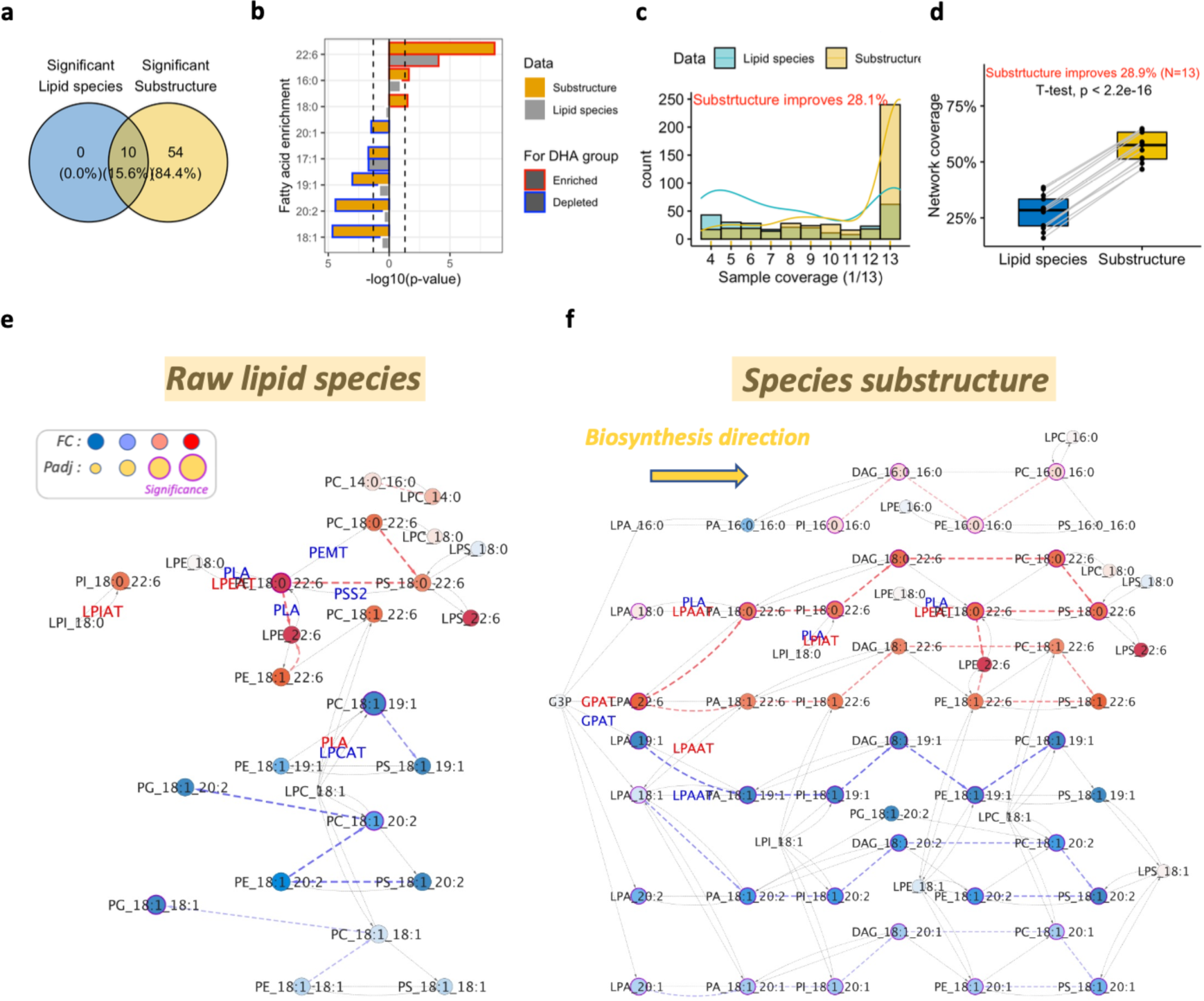
Substructure analysis improves the non-independence and sparsity of measured lipids, thus improving the statistical analysis of the differences across conditions and improving interpretability. **a**, A Venn diagram shows the increased number of significant features (lipids and substructures) with iLipidome (adjusted p-value<0.05) in DHA-treated RBL cells. **b**, Enrichment analysis for significant lipid species or substructures based on FA categories. **c**, Proportion of samples containing a lipid or substructure in 13 samples, and the associated probability distribution. **d**, The coverage in entire biosynthetic network for 13 lipid profiles using lipid species or substructures. The improvement index in (**c**) and (**d**) is calculated based on the average sample or network coverage from the raw lipid species and the iLipidome substructure data. **e**,**f**, The lipid biosynthetic network built from all significant pathways included in the top 3 activated and suppressed representative pathways in **Extended Data** Fig. 2d,e. Red and blue indicate activation and suppression while color depth denotes the importance of pathways. The top 5 perturbed edges in **Extended Data** Fig. 2f,g are also labeled in the network. GPAT, glycerophosphate acyltransferase; LPAAT, lysophosphatidic acid acyltransferase; LPCAT, lysophosphatidylcholine acyltransferase; LPEAT, lysophosphatidylethanolamine acyltransferase; LPIAT, lysophosphatidylinositol acyltransferase; PLA, phospholipase A; PEMT, phosphatidylethanolamine N-methyltransferase; PSS2, phosphatidylserine synthase 2.

### iLipidome reveals enzyme FA preference and suggests potential therapeutic targets for GBM

The Lands cycle is crucial for FA remodeling in membrane structural lipids, which can further impact membrane properties and functions^2,33,34^. Here, we applied iLipidome to uncover the hidden interactions by LPCAT1 in the Lands cycle and to explore promising therapeutic targets with a GBM dataset^2^ (**Fig. 5a**). In the dataset, their LC/MS lipidomics analysis could not differentiate each FA in phospholipids (e.g., PC 32:0 instead of PC 16:0-16:0). We found saturated PCs were most suppressed after LPCAT1 knockout but it is difficult to deduce their relationships (**Extended Data Fig. 3a**). We accounted for the interdependence among lipids and were able to increase statistical power and build the connections between lysophospholipids (one FA) and phospholipids (two FAs) (**Fig. 5b and Extended Data Fig. 3b**). LPCAT1 has specificity to synthesize saturated PC from saturated FAs and LPC^33,35^. As expected, the pathways involving saturated PC were suppressed in the LPCAT1 knockout group (**Fig. 5c**). Simultaneously, we found three saturated LPC-PC reactions were most inhibited in the Lands cycle in the substructure network (**Fig. 5d**). In contrast, no pathways showed significant and only two LPCAT1 reactions were identified when we analyzed the raw lipid species data (**Extended Data Fig. 3c,d**). The interactions unraveled by iLipidome allowed us to understand the exact chain length and double bond number of PCs (12:0-16:0, 14:0-16:0, and 16:0-16:0), along with free FAs involved (12:0, 14:0, and 16:0) (**Fig. 5b**). Our findings matched the enzyme activity experiment showing that these saturated FAs were three of the most preferred substrates for LPCAT1^35^. As lauric acid (12:0), myristic acid (14:0), and palmitic acid (16:0) are the major components of palm oil and coconut oil, the FA information exposed by iLipdiome also raises their dietary source as having potential relevance to GBM, alongside treatments modulating LPCAT1. Finally, a corresponding accumulation of free 16:0 FA was validated in the LPCAT1 knockout groups (**Fig. 5e**). Altogether, the unique function in iLipidome gives us an opportunity to dissect exact FA composition, interrogate enzyme substrate specificity, and identify dietary lipids associated with disease and candidate treatments.

**Fig 5:**
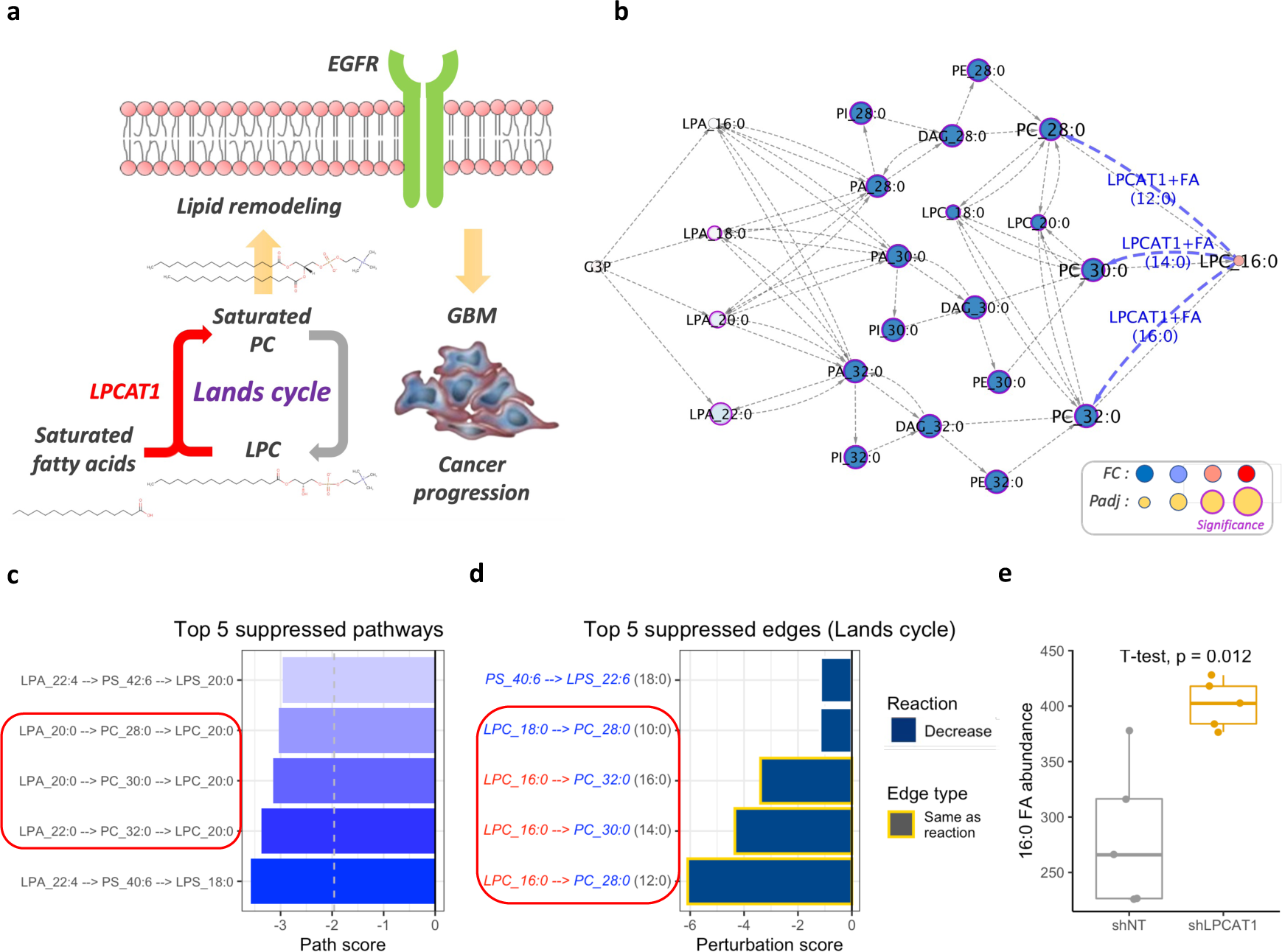
Identification of LPCAT1-associated reactions and its FA specificity by iLipidome lipids associated with GBM and potential treatments. **a**, LPCAT1-mediating membrane lipid remodeling is required for sustained EGFR signaling and promotes GBM cell growth. **b**, When comparing wild type and LPCAT1 knockout U87EGFRvIII GBM cells, the iLipidome substructure network built included three significantly suppressed pathways containing saturated PCs. **c**, Top 5 significantly suppressed pathways in the substructure analysis. Pathways are simplified and represented by the first and the last metabolite. If the last metabolite belongs to lysophospholipids, we add the second to last metabolite. **d**, Top 5 Lands cycle’s reactions that are significantly suppressed in the network built from (**c**) We highlighted the pathways or the reactions containing saturated PC with a red frame in (**c**) and (**d**). **e**, Boxplots display a significant increase in 16:0 free FA in the LPCAT1 knockout groups.

### iLipidome explores molecular mechanisms for cardiovascular diseases (CVDs) and facilitates early biomarker discovery

Disturbances in lipid homeostasis have been linked to many disorders, making lipidomics analysis a powerful tool to investigate disease mechanisms and to search for biomarkers^36–38^. Normally, most significant lipids will be selected for further analysis, but they can be limited to high variability in lipidomics data and barely explain the fluctuations of the entire lipidome. To expand the applicability of iLipidome for exploring mechanisms and biomarker candidates, we applied our method to a dataset composed of 146 plasma lipid profiles from 85 controls and 61 potential CVD patients (**Fig. 6a**)^4^. The 61 individuals went to the hospital with chest pain and displayed a similar lipidomic pattern with those having more severe CVD symptoms. This suggests that it may have both therapeutic and diagnostic value to analyze lipid alterations in these patients.

**Fig 6:**
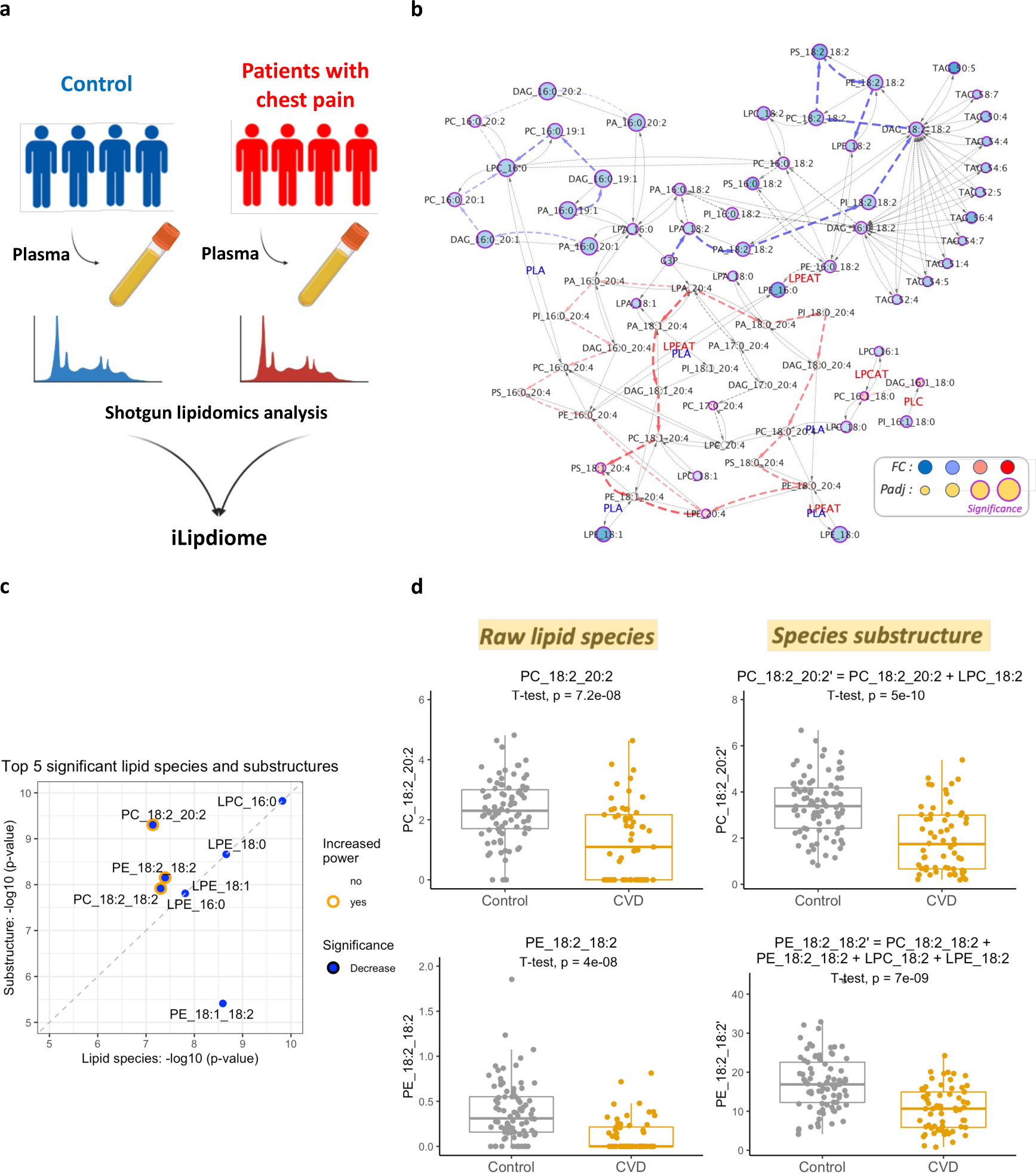
iLipidome unveils the lipid dysregulation and identifies early biomarker for cardiovascular diseases (CVDs). **a**, A schematic of the experiment comparing plasma lipidomics between controls and CVD patients. **b**, The substructure network built from all significant pathways included in the top 5 activated and suppressed representative pathways in **Extended Data** Fig. 4d. The top 5 perturbed edges in **Extended Data** Fig. 4e are also labeled in the network. **c**, Paired comparison of the statistics from top 5 significant raw lipid species and substructures. Only those lipids not missing in raw lipidomics data are considered. If the substructure method improves the statistical power, the lipids will be colored in orange. **d**, Boxplots show species substructures are more stable and have a higher discriminatory power than raw lipid species. LPCAT, lysophosphatidylcholine acyltransferase; LPEAT, lysophosphatidylethanolamine acyltransferase; PLA, phospholipase A; PLC, Phospholipase C.

First, iLipidome significantly improves the statistical power in this dataset (**Extended Data Fig. 4a**). The pathway analysis revealed that lipids with 18:2 FA and 20:4 FA were enriched in the suppressed and active pathways, respectively (**Fig. 6b and Extended Data Fig. 4b,c)**. The major 18:2 FA in human body is LA, which can normalize blood cholesterol and triglyceride level through proprotein convertase subtilisin/kexin type-9 (PCSK9)^39,40^ and negatively correlate with CVD risk^41^. Otherwise, AA is the main form of 20:4 FA and an important precursor of many inflammatory mediators, such as hydroxyeicosatetraenoic acid (HETE) and leukotrienes ^42^. Their critical roles in inflammation and coagulation may explain their positive association with CVD risk in our and the other studies^43–46^. Moreover, growing evidence indicates arachidonoyl phospholipids (PLs), especially PE, can act as substrates for lipid peroxidation that result in ferroptosis and cell death^47,48^. Our iLipidome analysis also showed the most altered reactions tend to accumulate AA in PC and PE in the CVD group (**Extended Data Fig. 4d,e**). Besides, most ferroptosis related reactions, such as the Lands cycle or LPAAT reactions involving AA addition, were consistently up-regulated (**Extended Data Fig. 4f,g**). As our substructure method can capture upstream intermediates LPA, it allows a unique chance to survey both types of reactions and their associated enzymes (**Extended Data Fig. 4g**). Since our lipidomics data came from plasma and these reactions usually occur in cells, it is difficult to determine where these AA-rich PLs were synthesized. However, their increase in plasma as free PLs or lipoprotein conjugates might have a higher chance to be incorporated into tissues through lipid transporters or lipoprotein receptors and exert related biological functions including inflammation and ferroptosis, both of which have been linked to CVDs^49–51^.

Our substructure method reveals potential mechanisms and summarizes most impacted lipid pathways, allowing us to find robust biomarkers for CVD. We compared the p-values from the top 5 significant raw lipid species and substructures (**Fig. 6c**). Although the most significant feature is LPC 16:0, lysophospholipids usually locate at the end of a pathway, so theoretically their statistics will not be affected by the substructure abundance of other lipids. On the contrary, we found the statistical power increases in three out of four phospholipids after transformation (**Fig. 6c**). Interestingly, these lipids all carry 18:2 FA, corresponding to the most suppressed pathways we identified (**Extended Data Fig. 4c**). A further paired analysis demonstrated that the substructures could aggregate lipid signals in a biosynthetic scenario and thus were more stable and showed a better discriminatory power than using raw lipid species alone (**Fig. 6d**). Moreover, iLipidome clearly points out the composite lipid units for each substructure, providing a more systems-based platform to optimize biomarkers. In summary, iLipidome allows intensive analysis of underlying mechanisms behind complicated lipidomics data and simultaneously guides robust biomarker discovery that can be translated into clinical application.

## Discussion

Lipidomic data are typically studied as if each lipid is independent^48,52,53^. iLipidome, in contrast, uses a data-driven approach combining FA information to search substructures for a lipid profile based on their biosynthetic pathways. These innovative features of iLipidome enrich small changes in shared lipid intermediates, resulting in statistically significant substructures and pathways. Firstly, we showed here with three example datasets that iLipidome elucidated significant increases in statistical power (**Fig. 3c and Extended Data Fig. 2a, 3b and 4a)** and better sample coverage (**Fig. 4c and Extended Data Fig. 1e**). Accounting for interdependence and interconnections enabled us to explore altered biosynthetic fluxes in a unified biological theme (**Fig. 3b, 4f, 5b and 6b**). Moreover, the advantages are also apparent in the further analysis, such as hierarchal clustering and enrichment. Substructure clustering displays a more biologically meaningful grouping of both the experimental conditions and the lipid features (**Fig. 3f,g and Extended Data Fig. 2e,f**). FA enrichment analysis for substructures also reveals a significant DHA-induced FA remodeling that may be missed in conventional analysis of lipidomics data (**Fig. 4b**).

As lipidomics data lack information about the lipid biosynthesis and interdependence, it remains difficult to pinpoint genetic causes of alterations in lipid profiles. Through iLipidome, we can now more easily find the enzymes and perturbed reactions that drive lipidome reprogramming across conditions. For example, the inhibitory effect of DHA or PUFAs on SCD1 has been extensively studied^3,13,26,54^. Our method successfully elucidated this relationship with the FA substructure data (**Fig. 3e and Extended Data Fig. 1b**). Moreover, iLipidome successfully attributed the increase of EPA following DHA to the inhibition of ELOVL2/5 and EPA elongation to docosapentaenoic acid (DPA, 22:5 omega-3) (**Fig. 3b**). Our results are further supported by the experiments showing that ELOVL2/5 are down-regulated by DHA and dietary ALA is necessary for the EPA accumulation with DHA feeding^28,29,55^. Interestingly, numerous desaturases and elongases were also identified to be affected in the DHA-treated samples (**Fig. 3e**). We note that iLipidome focuses on the biosynthetic pathways for omega-3 and omega-6 fatty acids and their intermediates, which may not be detected in the lipidomics data due to complete conversion to larger lipids or limitations of the detection platform. However, iLipidome enables the exploration of likely changes in FA biosynthetic pathways and reactions from the limited data available. Thus, while it is crucial to interpret some results in sparsely measured pathways with caution, it provides a mechanistic way to reliably estimate likely pathway activities. It will be important for users to validate their specific results and follow up with additional experiments. For example, it would be valuable to clarify whether common regulators of lipid metabolism, such as sterol regulatory element-binding protein (SREBP) or proliferator-activated receptor (PPAR), have a role to play in the response to DHA supplementation^32,56^. We found the genetic source to induce FA reprogramming after DHA supplementation and saw the effect can be profoundly passed to all lipid species (**Fig. 3b and 4f**). Also, our substructure decomposition can capture lipid intermediates and enables one to explore the upstream reactions (e.g., LPAAT) that could induce wide-ranging phospholipid remodeling (**Extended Data Fig. 2j**). In the other dataset, we identified the reactions to accumulate arachidonic PE and PC increased in the CVD patients (**Extended Data Fig. 4c**). These plasma lipids could potentially induce inflammation or ferroptosis and contribute to the development of CVD^43,45,49,50^. Since the reactions involving AA addition primarily occur in cells, it might be valuable to investigate where these reactions and associated enzymes (e.g., LPCAT3 and AGPAT3) involving AA transfer are dysregulated in the early stage of CVDs. LPCAT3 has LPEAT and LPCAT activity, while AGPAT3 (LPAAT) catalyzes LPA reactions^33^. To further examine the general applicability of our method, a dataset containing multiple gene knockouts in lipid metabolic pathways was also used to assess the performance of iLipdiome^57^. Overall, the top 5 perturbed pathways across 11 gene knockouts revealed by iLipidome were highly consistent with the enzyme specificities and the expected lipidomic changes affected by each knockout (**Extended Data Table 1**). More essentially, five out of six genes that appear in our reference pathway were also ranked in the first place of their respective most suppressed reactions. Taken together, iLipidome sheds light on the genetic source of complex lipid profiles. These insights will guide future research into the mechanisms regulating lipid disorders or interactions between PUFAs and endogenous FA biosynthesis.

Another strength of iLipidome is its ability to reconstruct the active biosynthetic network in measured samples. Indeed, the low concentrations of lipid intermediates make it hard to link reactants and products in a lipid profile^33,58^. Compared to BioPAN^10^ (**Extended Data Fig. 5a-c**), our substructure method forms a network with a high degree of coverage and connectivity (**Fig. 4d and Extended Data Fig. 1f**). It might be because the BioPAN algorithm requires free FA data to connect the reactions involving FA transfer. In addition, iLipidome offers a more comprehensive analysis pipeline that uses the constructed network to identify lipid intermediates/reactions and improve statistical power and data sparsity in contrast to other lipid network analysis tools (**Extended Data Table 2**)^59^. To compare the results obtained from three existing tools, we conducted an analysis using a new dataset with free FA data to explore the impact of dietary restriction (DR) on lipidomic changes^60^. The study revealed that mice subjected to dietary restriction exhibited significant lipid reprogramming, with a notable increase in the synthesis of phospholipids, particularly PC and cardiolipin (CL). In addition, pathways involved in the degradation of membrane lipids or their conversion to triacylglycerol (TAG) were significantly less active. iLipidome accurately identified the comprehensive alterations in the lipidome while BioPAN and LINEX2 only captured partial information regarding PC and TAG changes (**Extended Data Fig. 6a-c**). Both iLipidome and BioPAN analyses revealed an obvious upregulation in de novo FA synthesis, desaturation, and elongation in young dietary restriction (DR) mice (**Extended Data Fig. 7a-c**), consistent with the transcriptional upregulation of de novo lipogenesis previously observed^60^. One of the key advantages of iLipidome is that it can extract FA residues from glycerophospholipids and glycerolipids as alternative sources to build the network in cases where free FA data is not available. The results obtained using FA residues were similar to those obtained using fatty acyls, as supported by the significant positive correlation observed between the expression levels of fatty acyls and FA residues (**Extended Data Fig. 7b,d**). This supports the robustness and reliability of iLipidome in capturing the FA network even in the absence of direct free FA measurements.

In iLipidome, we have two unique systems to decompose lipids according to different structural resolutions. First, FAs should be inherited in lipid species using the same biosynthetic pathways. Second, for those lipids lacking the exact identity of FAs (e.g., PC 30:0)^2,48^, they will be matched to the LIPID MAPS Structure Database (LMSD)^23^ and find possible FA combinations. These methods allow us to maintain the completeness of biosynthetic networks and the important FA information, thus extending the method’s applicability in searching potential therapeutic targets and clinical biomarkers. For example, high saturated PC levels produced by LPCAT1 are required to maintain EGFR signaling and GBM cell growth^2^. Our algorithm successfully established clear links between LPC 16:0 and three saturated PC (28:0, 30:0, and 32:0) (**Fig. 5b**) and suggested two compelling therapeutic strategies for GBM: (1) inhibition of LPCAT1 and (2) depletion of free FAs, especially lauric acid (12:0), myristic acid (14:0), and palmitic acid (16:0). iLipidome not only filled the gap of FA information but also correctly explained the substrate specificity of LPCAT1^35^. In the CVD dataset, the inherited feature of lipids sharing the same FAs helped connect each substructure and can be applied to identify novel lipidomic biomarkers. By decoding substructures, we clearly see how lipid signals aggregate in a biosynthetic pathway (**Fig. 6d**). This process allows iLipidome to explain interdependence among measured lipids, thereby improving statistical power and reducing sparsity, thus making substructures a more stable and precise biomarker. Altogether, this functionality will be invaluable for interrogating the specificity of the acyltransferase family members and its effect on the membrane protein behavior. Furthermore, it provides an ideal platform for researchers to design reliable lipid biomarkers in a systems biology scenario.

## Materials and Methods

### Demonstrating lipidomics datasets

We demonstrate the developed functionalities of iLipidome on four publicly available lipidomics datasets^2–4,57^. Except the CVD dataset, we adopted the same data-processing approach. All of our datasets used internal standards to quantify measured lipids and obtained their picomole values (absolute quantification). First, we excluded the lipid species with missing values in over 70% samples to reduce the noise. The default parameter for this step in MetaboAnalyst^61^ and LipidSig^5^ is 50%, which we used in the larger CVD dataset (146 samples). On the contrary, since the other three datasets have fewer samples (13, 10, and 6 or 9 samples for each comparison), we relaxed the threshold (70%) to include more lipid species and to better reflect comprehensive lipidomic changes. Secondly, the values were transformed into mol% to obtain a final lipid profile. The first dataset explored the lipidomic and biophysical homeostasis of mammalian membranes as influenced by dietary lipids, which contains 13 membrane lipid profiles including 7 controls and 6 DHA treated rat basophilic leukemia (RBL) cells^3^. To investigate membrane lipid metabolism, 444 individual lipid species across 16 lipid classes in GPLs and diglycerol (DAGs) were selected for the following analysis (**Supplementary Table 1**). We also computed the FA profiles through the decomposition of lipid species into FAs and the summation of their values (**Supplementary Table 2**). Two different kinds of profiles were used to calculate FA or species substructures, respectively. We removed ether lipids when computing species substructures since we could not differentiate alkyl and alkenyl ether lipids. It might introduce a bias if we directly map them to our reference pathways, where two kinds of ether lipids are separated. Further, the biosynthetic pathways of ether and ester lipids are bifurcated at glycerol-3-phosphate, the start point of glycerophospholipid biosynthesis. Hence, the backpropagation process of ether lipids would have a minor effect on the results we obtained from ester lipids only. The second dataset studied the role of LPCAT1 in the transduction of oncogenic signals through shaping membrane lipid composition, which contains 10 lipid profiles of human glioblastoma cell line (U87EGFRvIII), expressing LPCAT1 shRNA or a non-targeting control^2^. There are 82 lipid species across 6 lipid classes and 4 free FAs (16:0, 18:1, 18:0, and 20:4) in this dataset (**Supplementary Table 3**). The abundances of lipid species were pre-processed using the common method above while free FAs were analyzed with raw values. For the CVD dataset, we collected 146 plasma lipid profiles including 85 controls and 61 potential CVD patients to explore the mechanisms and early biomarkers for CVDs^4^. Those patients went to the hospital with chest pain but without indicators for stable angina pectoris, unstable angina pectoris, or myocardial infarction. The lipid species missing in over 50% samples were excluded, resulting in an expression table composed of 197 lipid species across 9 lipid classes (**Supplementary Table 4**). To reflect the actual concentration of lipids in the plasma, we adopted the raw lipid values to analyze this dataset and design the biomarkers. The fourth dataset investigated multiple gene knockouts in lipid metabolic pathways and explored how they affected the lipidome in the human colorectal carcinoma (HCT116) cell line^57^. We selected the lipid profiles of 2 control groups based on the different knockout method (gene trap or deletion) and of 11 gene knockout groups whose effects on the lipidome have been clearly described in the paper. Each group contains at least three replicates. In total, 30 lipid profiles (3 controls and 27 knockouts across 8 genes) from the gene trap method and 15 lipid profiles (3 controls and 12 knockouts across 3 genes) from the deletion method were collected for the following analysis. Two methods also result in 1836 and 1923 lipid species across 24 lipid classes, respectively (**Supplementary Table 5**). The final dataset aimed to explore the effects of dietary restriction on lifespan extension and metabolic health^60^. The lipidomics data of white adipose tissues was collected from four mice fed ad libitum (AL) and four mice subjected to dietary restriction (DR) in their early age (**Supplementary Table 6**). The dataset comprises 516 lipid species belonging to 28 lipid classes and three classes of fatty acyls, including free fatty acids (FFA), fatty acyl CoAs (FaCoA), and fatty acyl carnitines (FaCN). In the subsequent FA analysis, we examined both the fatty acyls and the FA residues extracted from major lipid classes and compared the results.

### Substructure decomposition based on biosynthetic pathways

#### Curation of the reference lipid biosynthetic pathways

To decompose lipids into substructures, FA and lipid biosynthetic pathways for human, mouse, and rat were obtained from two widely used pathway databases (KEGG^15^ and Reactome^16^). In this study, we collected 6 KEGG pathways (Fatty acid biosynthesis, Glycerolipid metabolism, Glycerophospholipid metabolism, Ether lipid metabolism, Sphingolipid metabolism, and Biosynthesis of unsaturated fatty acids) and 6 Reactome pathways (Fatty acyl-CoA biosynthesis, alpha-linolenic (omega3) and linoleic (omega6) acid metabolism, Triglyceride metabolism, Phospholipid metabolism, Sphingolipid metabolism, and Wax and plasmalogen biosynthesis) to cover most common FAs and lipid classes in lipidomics analysis^1–4^. Together with recent studies and reviews on the metabolism of polyunsaturated FAs^17–19^, ether lipids^20,21^, and deoxysphingolipids^22^, we manually curated the resulting reference lipid biosynthetic pathways, which compose of 64 lipid class (**Supplementary Table 7**) and 49 FA related nodes (**Supplementary Table 8**). Currently, the reference pathways in iLipdiome cover generic biosynthetic routes in human, mouse, and rat and not tissue-specific or cell type-specific lipid biosynthetic routes. For each target lipid, we search all possible routes to synthesize it with the *all_simple_paths()* function of R igraph package. The metabolites in the routes are considered as substructure candidates. Note that the source vertex differs according to the FA or lipid class metabolism pathway each lipid belongs to (e.g., G3P for GPLs, glycerolipids, and ether lipids; serine and palmitoyl-CoA for sphingolipids; alanine and palmitoyl-CoA for deoxysphingolipids; acetyl-CoA, linoleic acid (omega-6), and alpha-linolenic acid (omega-3) for all FAs).

#### Fatty acid substructure decomposition

FA decomposition generally follows three major biosynthetic pathways: non-essential FA biosynthesis, omega-6 FA biosynthesis, and omega-3 FA biosynthesis. Since some of our lipidomics datasets don’t provide exact double bond position, a FA could be mapped to multiple targets (e.g., 18:3 for omega-6 and omega-3 FAs). For these targets, the expression is generally equally distributed and the pathway search will be done separately. Among them, 18:2 FA and 20:4 FA cross different biosynthetic pathways and account for a large proportion in total FA as LA (18:2 omega-6) and AA (20:4 omega-6). Since they will have a huge impact in substructure computation, we identify them as their major form instead of equal distribution. Besides, palmitate (16:0) is the first FA produced in de novo FA biosynthesis^62^. The FAs shorter than it may be the intermediates or come from exogenous sources. Therefore, we separate them to endogenous and exogenous pathways in the beginning. Only those in the endogenous pathways will be further decomposed. Finally, in the DHA dataset, we did not process DHA and its neighbor FAs (22:5 and 24:6 omega-3) to control the artifacts owing to supplementation.

#### Lipid class and species substructure decomposition

The iLipidome algorithm can decompose lipid species into lipid class and species substructures based on the lipid pathways we collected. For the lipid class substructure, our reference lipid biosynthetic network composed of lipid classes allows us to directly decompose them. The lipid species substructures, on the other hand, further require the FA information encoded in lipid species and the LIPID MAPS database^23^ to decompose. Thus, substructures can represent precursors or intermediates in potential biosynthetic routes to synthesize certain lipid species and be used to reconstruct the following lipid molecular network. There are four major steps for processing FA information of the lipid species substructures from lipid biosynthetic pathways. Firstly, if the substructures in one pathway have the same FA number as the target species (e.g., DAG in PC pathway and both of them have two FAs), we assigned the same FA composition of the target to all these substructures (e.g., DAG 16:0-18:1 for PC 16:0-18:1). Secondly, for those substructures *S* with less FA number (e.g., LPA in PC pathway but it only has one FA), we find the lipid class *k* closest to the target species and list all possible FA combinations of it *S_k_*_1_,_*k*_2_,…,_*_k__n_* based on the target (LPA 16:0 and LPA 18:1 for PC 16:0-18:1). Only the substructures *S_k_*_1,_*_k_*_2,…,_*_k__n_* appearing in lipidomics data will be kept, which allows us to control the number of potential substructures. If none of the substructures match, we retain all of them, each of which represents one biosynthetic route. Besides, the FA combinations defined above will apply to the other substructures in *S* with the same FA number. Third, for the substructures with more FA number (e.g., PC in LPC pathway), we also find the closest lipid class and select those with the same FA as the target species in lipidomics data (PC 16:0-18:1 for LPC 16:0). If there is no match, we will remove the FA information of the substructure (PC for LPC 16:0). Fourth, some datasets might not provide the exact FA identity for their lipids (e.g., PC 34:1 instead of PC 16:0-18:1), which will cause difficulties in processing the substructures with different FA number. To address this problem, target lipid species with unknown FA composition are matched to the LIPID MAPS Structure Database (LMSD)^23^ to find all possible FA combinations of them (PC 16:0-18:1, PC 16:1-18:0, etc.). With the FA information, we can decompose these lipids via the approach mentioned above. We note that the imputation of fatty acyl compositions for the target species is only performed to identify FA information for the substructures with different FA number (e.g., LPA 16:0 and LPA 18:1 for PC 34:1). The substructures with same FA number will still remain the original FA form as PC 34:1 and DAG 34:1. Furthermore, if the FA composition of the closest substructure is unknown in lipidomics data (e.g., how to find PC 34:1 in LPC 16:0 pathway), we will search the candidate species with the same lipid class as the substructures (PC) and the same FA as the target species (16:0) using the LMSD database. The FA information of candidate species in the database can be summed (e.g., PC 16:0-16:1 and PC 16:0-18:1 to PC 32:1 and PC 34:1) and used to screen in the lipid profiles so that we have more confidence to build the connections between LPC 16:0 and PC 32:1 or PC 34:1. Finally, we do not consider the exact position of FAs in relation to the glycerol backbone (sn1 or sn2) except for the ether lipids. To summarize, these processes enable us to precisely identify the substructures and connect biosynthetic pathways using available information in a lipid profile and pave the way for subsequent analyses.

### Substructure extraction and computing substructure profiles

To identify the potential source of synthesized lipids and correct the impact of exogenous lipid uptake, the substructures in each route were mapped with their fold change from raw lipidomics data and were extracted through a backpropagated process. From the target FA or lipid species, the checking process along the biosynthetic route would not stop until it meets a substructure with an opposite fold change. One exception is the endogenous biosynthesis pathway for FAs in the upstream of palmitate. Because they are synthesized as a group, we do not check their fold change and keep all substructures. Only those passing the examination are selected with their weights initialized as 1. If there is more than one biosynthetic route for a FA or lipid species, the weight is equally divided and distributed to each path. Because we allow the substructures that did not appear in our dataset to pass the backpropagation, a threshold to control the percentage of missing data in one pathway is set to reduce artifacts. Typically, the threshold from 70% to 100% can be adjusted according to total lipid class number in each dataset. If the proportion of missing substructures exceeds the threshold in one biosynthetic route, the target FA or lipid species will not be decomposed to substructures (**Extended Data Fig. 8**). The checking process of fold changes and the threshold of missing substructures can prevent unlimited backpropagation of substructures and over-weighting certain pathways. To construct a substructure matrix, the substructures with their weights from each FA or lipid species are matched to a list composed of all substructures extracted from a dataset. A substructure matrix encoding the frequency of each substructure is then multiplied by the lipid profile to produce the resulting substructure profile, which records the abundance of each substructure and can be used to reconstruct the biosynthetic network. We note that substructure profiles are calculated using a direct add-up function for selected substructures regardless of data source. Although similar method has been widely used to analyze lipid features, such as double bond and chain length, in many studies^3,34,63^, users should consider whether their data are absolute quantification and suitable for this process given that different lipid classes have different ionization tendencies depending on their headgroup.

### Pathway scoring method

For any two nodes in the biosynthetic network, we search all possible pathways using the *all_simple_paths()* function from the R igraph package. We then explore the active and suppressed ones in the pathway set using the method adapted from^64^. Specifically, we transform the p-value *p_i_* of each node *i* in a pathway into z-score, which will be used to calculate the pathway score. More specifically, given that t-distribution can be approximated by a normal distribution, each p-value is converted to a z-score through 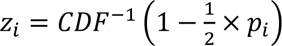, where CDF is the cumulative distribution function. If the fold change between two experimental conditions is lower than 1, we add a minus sign to *z_i_*. For a random data, *z_i_* will follow a standard normal distribution. To calculate the score of a pathway *A* of *n* nodes, we summed all *z_i_* in the pathway by taking 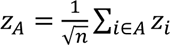. Because the variance of a sum is the sum of the variances for independent random variables, *z_A_* will also follow a standard normal distribution if the *z_i_* are independently drawn from a standard normal distribution. A high *z_A_* indicates an active pathway while a low *z_A_* corresponds to a suppressed pathway. In addition, the length (n) of pathway is considered in the function helps us not over-weight certain pathways. Further, to calibrate *z_A_* against the background distribution, we adopt a Monte Carlo approach to randomly select *n* nodes and compute their scores 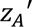. The values are used to derive the mean 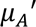 and standard deviation 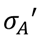 for each *n*. *z_A_* is corrected by these estimates to produce a final path score 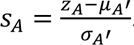. With this calibration, we can reduce noise and guarantee to have µ = 0 and σ = 1 for a randomized dataset. To determine if a pathway is significantly activated or suppressed, we choose the critical value 1.96 as the threshold based on the significant level (p-value<0.05) from a two-tailed test. Consequently, *S_A_* > 1.96 and *S_A_* < −1.96 represent the significantly activated and suppressed pathways, respectively.

### Identify representative pathways

For all pathways found in the biosynthetic network, we use two methods according to FA or lipid species to classify them into different representative pathways. At first, pathways are separated to up-regulation or down-regulation based on the signs of their scores. We then rank their impotence using the absolute value of path scores, followed by a pathway similarity search from top to bottom. The pathway with the largest score is assigned as the first representative pathway. In the FA analysis, we sequentially compute the proportion of overlapping edges between the candidate pathway and the representative pathway set. If the overlapping proportion of a pathway is over 50% with one representative pathway, it will be classified into that one. If there are no representative pathways over 50%, the candidate pathway will be a new representative pathway. On the other hand, because the FA information is inherited in the lipid species through biosynthetic connections, we can find the dominate FA composition with highest frequency (e.g., 16:0-18:1 or 32:0) in each lipid species pathway. Similar to the FA analysis, pathways with the same dominate FA will be categorized to the same representative pathway. If the dominate FA is the first time appear in this process, the pathway will be considered as a new representative pathway. In this study, we typically collected all significant pathways belonging to the top 3 or top 5 representative pathways for constructing the species biosynthetic network.

### Reaction scoring

To determine the potential driver reactions or enzymes in the biosynthetic network, we developed a novel algorithm to compute the perturbation score for each reaction (edge). The concept is to compare the ratio of product over reactant and evaluate its statistical significance magnitude across conditions. Firstly, we can compute this ratio in a certain reaction for each sample. Then, the fold change is obtained by the averaged ratio from control group divided by the averaged ratio from experimental group. Next, we use two-tailed Student’s t-tests to compute the p-value of ratios between different conditions. With these values, the final perturbation score of reaction is calculated by: −*log*_10_(*p* − *value*) × *log*_2_(*fold change*). The score is considered significant when the p-value lower than 0.05. Genes/Enzymes involved in reactions are labelled based on the KEGG^15^ and LIPID MAPS^23^ database and other recent studies^17–19,22^. Note that since these databases and studies only provide information at the lipid class level, the corresponding genes/enzymes in lipid molecular reactions highlighted by iLipidome do not consider lipid species specificities. In our figures, we only chose one of them to represent the reactions.

### Hierarchical clustering

Clustering analysis and heatmap visualization was done in R with the package ComplexHeatmap^65^. Each lipid variable was scaled to have mean 0 and standard deviation 1 before clustering. The distance matrix, defined as 1 minus the Spearman correlation, was processed by the complete method. Further, the *cutree()* function was applied to split the samples and FA variables into two groups. To compare the similarity between the clustering results and the original labels, we calculated the adjusted Rand index using the R package GeometricMorphometricsMix^66,67^, and the statistical significance is further tested by permutation test. FA variables were categorized according to the double bond number with saturated FAs and highly unsaturated FAs (3 or more double bonds) in one group and less unsaturated FAs (1 and 2 double bonds) in the other.

### Enrichment analysis

We performed two different enrichment methods to explore and compare fatty acid signature between substructure and unprocessed data. First, lipid species or substructures were separated into increased and decreased groups according to fold change in advance. For each group, we further categorized them by significance (P-value < 0.05) and FA composition. Finally, enrichment analysis for each FA category was performed using over representation analysis with Fisher’s exact test. On the other hand, all lipid species or substructures were only ranked according to their statistics and grouped by FA composition. An enrichment analysis was further performed using gene set enrichment analysis (GSEA) in the ‘*fgsea*’ package^68^.

### Statistical Analysis

All statistical analyses were performed on R (v 4.1.0) platform. Two-tail Student’s t tests or moderated t-tests in the limma package^69^ were used for comparing two experimental groups. We applied Benjamini–Hochberg correction to compute the adjusted p-values for a multiple testing. Statistical methods, numbers of samples, and p-values for each section are also indicated in the figure legends or methods.

## Data availability

All lipidome data used in this study were downloaded from the publicly available datasets and have been deposited with their code in GitHub (https://github.com/LewisLabUCSD/iLipidome). Source data for each figure/table are also provided with this paper.

## Code availability

We developed an R-Shiny web-based interface (http://www.bioinfomics.org/iLipidome/) that allows for user-friendly and interactive data analysis. In addition, we have packaged the code into an R package (https://github.com/LewisLabUCSD/iLipidome-package). It offers advanced users enhanced flexibility, accompanied by detailed documentation and example code for their convenience and reference. R files and functions to perform iLipidome analysis and generate all figures in this study are also available through GitHub (https://github.com/LewisLabUCSD/iLipidome).

## Acknowledgements

This work was supported by NIGMS (R35 GM119850) and generous support from the Novo Nordisk Foundation provided to the National Biologics Facility at the Technical University of Denmark (NNF20SA0066621). It was also supported by National Health Research Institute: grant # NHRI-EX111-11110BI, National Science and Technology Council: grant # MOST 111-2320-B-039-011, and China Medical University Hospital: grant # DMR-110-231, and # DMR-111-118. WJL was supported by the Elite Cultivation Scholarship Program Sponsored by China Medical University.

