## Supplementary Information for "iLipidome: enhancing statistical power and interpretability using hidden biosynthetic interdependencies in the lipidome"

#### Supplementary Text

**Text S1. iLipidome improves the non-independence and sparsity of measured lipids and increases the interpretability of lipidomic data by reconstructing its biosynthesis**

#### Supplementary Tables

**Table S1. Dataset 1 – Lipid species profiles**

**Table S2. Dataset 1 – Fatty acid profiles**

**Table S3. Dataset 2 – Lipid species profiles**

**Table S4. Dataset 3 – Lipid species profiles**

**Table S5. Dataset 4 – Lipid species profiles**

**Table S6. Dataset 5 – Lipid species and fatty acyls profiles**

**Table S7. The resulting references of lipid classes and lipid biosynthetic pathways**

**Table S8. The resulting reference fatty acids and fatty acid biosynthetic pathways**

#### Extended data figures

**Extended Data Fig. 1. Raw FA and substructure analysis for the DHA-treated dataset.**

**Extended Data Fig. 2. Raw lipid species and substructure analysis for the DHA-treated dataset.**

**Extended Data Fig. 3. Raw lipid species and substructure analysis for the LPCAT1 knockout dataset.**

**Extended Data Fig. 4. Raw lipid species and substructure analysis for the CVD dataset.**

**Extended Data Fig. 5. Application of BIOPAN's method in our datasets.**

**Extended Data Fig. 6. Comparison of the results from three tools for the dietary restriction dataset**

**Extended Data Fig. 7. Comparison of the FA network from iLipidome and BIOPAN for the dietary restriction dataset.**

**Extended Data Fig. 8. A workflow demonstrating how iLipidome manages substructure decomposition for the target lipids.**

#### Extended data tables

**Extended Data Table 1. The overall performance of iLipidome in identifying the essential pathways and reasons using a multiple gene knockout dataset.**

**Extended Data Table 2. Comparison of tools for the network analysis of lipidomics data.**

#### Text S1. iLipidome improves the non-independence and sparsity of measured lipids and increases the interpretability of lipidomic data by reconstructing its biosynthesis

Here, we decompose lipids from the DHA datasets into lipid species substructures and reconstruct biosynthetic network. Surprisingly, our substructure method gained additional statistical power and explored around five times more significant substructures than non-transformed lipid species (64 vs. 10) in the DHA-treated dataset (**Fig. 4a**). Moreover, enrichment analysis for significant substructures revealed a comprehensive FA remodeling consistent with our previous conclusion in FA substructure analysis (increase in 16:0 FA and 18:0 and decrease in 18:1 FA, 20:1 FA and 20:2 FA) (**Fig. 4b**). On the contrary, analysis for raw species data still failed to capture this FA signature. Considering that the missing values or statistical test methods might impact our results, we imputed the missing values with 1/2 of the minimum of each variable and computed the p-values using the moderated t-test. Consistently, iLipidome remarkably increased the statistical power and the number of significant species (68 vs. 18) and enhanced the following enrichment analysis (**Extended Data Fig. 2b-d**). Further, our lipid set enrichment analysis (LSEA) for the substructures also showed a higher normalized enrichment score (NES) and lower enrichment p-values than the unprocessed data (**Extended Data Fig. 2e**).

The increased statistical power in substructures may be due to the improvement of overall data quality. Accordingly, we found iLipidome mitigated statistical challenges by decreasing the proportion of lipids with a very low coverage across samples (**Fig. 4c**). Besides, correcting for sparsity also improves the hierarchical clustering in the scope of lipid species. The dependent species substructures display a higher adjusted Rand index value and a more consistent clustering on the DHA samples than the non-dependent lipid individuals (-0.032 to 0.435) (**Extended Data Fig. 2f,g**). On the other hand, our substructure-based method enhances the integrity of the biosynthetic network by raising the coverage up to 30 percent (**Fig. 4d**).

We further investigated whether interpretability of lipidomic data can be increased by iLipidome, specifically through analysis of the lipid substructures, visualized in the context of the biosynthetic network. Our results show that iLipidome can present the lipidomic data in a more complete and informative network (**Fig. 4e,f**). It allows us to clearly see how a phospholipid is synthesized from its very basic unit, glycerol-3-phosphate (G3P). This is especially valuable for datasets where there were perturbations upstream in lipid biosynthesis, since quantification of undetectable intermediates could prove more informative about the cause of the changes in the lipidome (**Fig. 4e,f**). To examine the robustness of iLipidome in searching critical pathways and reactions, a comprehensive comparison between the network built from raw lipid species and substructures was performed. First, we introduced our scoring system to measure pathway activities. In contrast to unprocessed data, the pathways in the substructure network are more significant and captured two major characteristics inferred from FA substructures: production of the lipids with omega-3 and saturated FAs and suppression of those with the other unsaturated FAs (**Extended Data Fig. 2g,h**).

To further investigate the mechanism behind them, we calculated and ranked the perturbed reactions and their associated enzymes (**Extended Data Fig. 2i,j**). The most impacted reactions are, as expected, strongly associated with the DHA incorporation. Furthermore, most belong to the Lands cycle or lysophosphatidic acid acyltransferase (LPAAT) catalyzed reactions, two major steps to generate FA diversity in GPLs[25]. This phenomenon is much more obvious when using the substructure method. The enzymes generally fall into three categories: (1) glycerophosphate acyltransferase (GPAT), (2) acylglycerophosphate acyltransferase (AGPAT) and membrane-bound O-acyltransferases (MBOAT) including LPAAT, LPCAT, lysophosphatidylethanolamine acyltransferase (LPEAT), and lysophosphatidylinositol acyltransferase (LPIAT), and (3) phospholipase A (PLA). The reactions controlled by AGPAT or MBOAT add one FA to lipids and are essentially the reverse reactions of PLA[25] (**Fig. 4e,f**). Considering DHA treatment in our dataset, the increased FA incorporation by AGPAT or MBOAT is more favored than the inhibition of PLA. Among them, LPAAT catalyzed reactions are the second step in the GPL biosynthesis and have potential to induce wide-ranging lipid reprogramming. As an intermediate, LPA is usually hard to detect in cells, making it impossible to investigate its reactions using raw lipidomics data[25] (**Fig. 4e**). However, our substructure method was able to address this problem and identified many LPAAT related reactions (**Fig. 4f**). On the other hand, the Lands cycle enzyme for DHA incorporation, such as LPEAT and LPIAT, were captured by both methods (**Extended Data Fig. 2i,j**). In previous research, LPAAT3 (AGPAT3)[59] and LPEAT2 (AGPAT7)[60] has shown high preference for DHA, making them potential drivers in the FA remodeling process. Yet, we cannot exclude the possibility that the rates of the reactions are regulated by DHA concentration instead of enzyme activity. On the other hand, GPATs transfer one FA to G3P and form LPA but they seem to play a less important role than the AGPATs or MBOATs since most GPATs exhibit the selectivity for saturated FAs instead of DHA[25]. We noticed a substantial suppression in phosphatidylethanolamine N-methyltransferase (PEMT) when analyzing raw lipid species data (**Extended Data Fig. 2i**). However, it contradicts the experiment showing that PEMT plays a role in synthesizing PC from PE enriched in DHA after DHA supplementation[61]. To summarize, iLipidome not only solves the interdependence issue to increase the statistical power but also improve the overall data quality by increasing the sample and network coverage. It allows for more reliable clustering and lipid enrichment analysis and more precise comparison of lipid biosynthetic pathways across conditions.

### Extended data figures

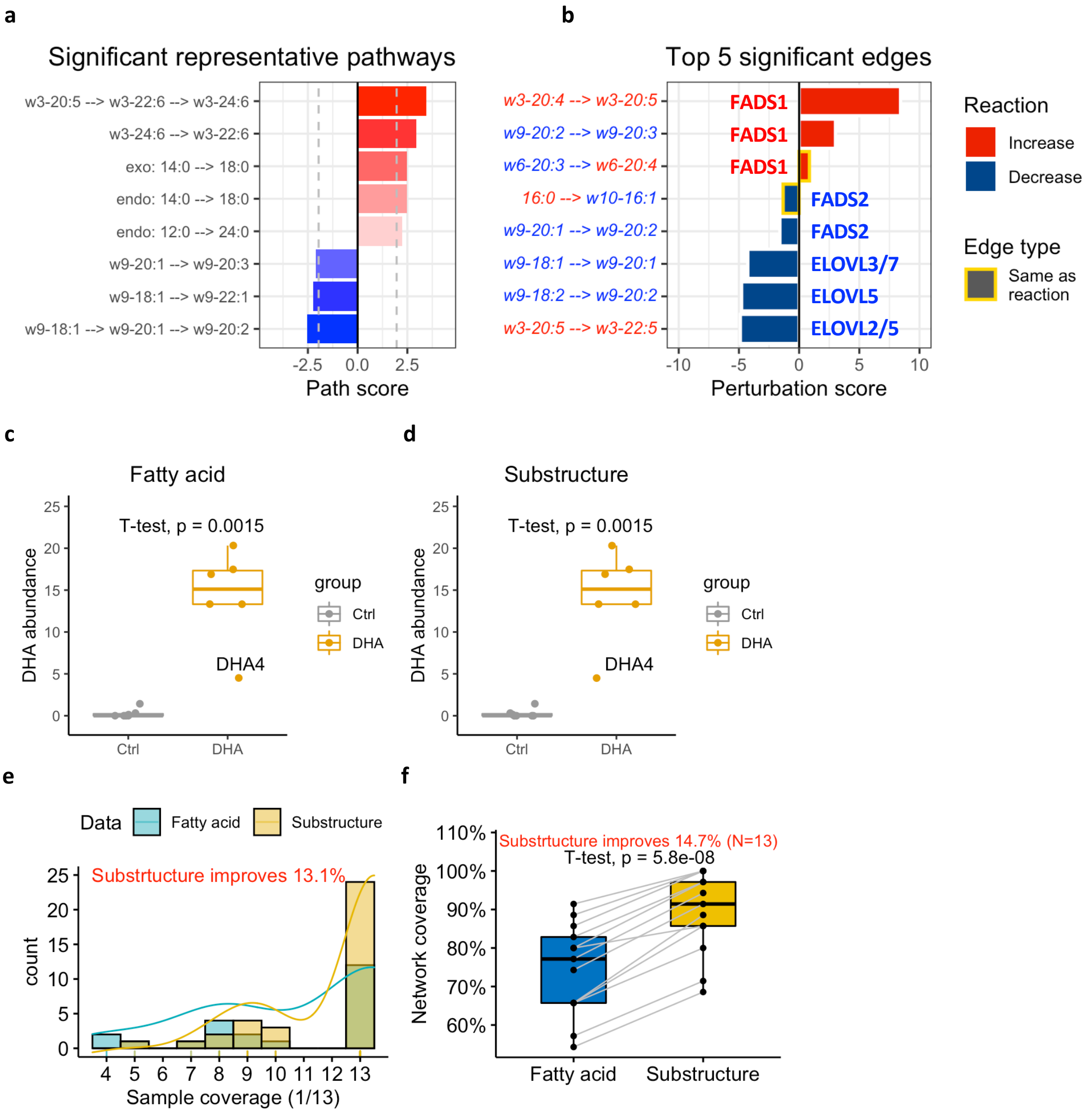


**Extended Data Fig. 1** Raw FA and substructure analysis for the DHA-treated dataset. **a**, Significantly activated and suppressed representative pathways in the network built from raw FA data. **b**, Top 5 significantly up-regulated and down-regulated edges (reactions) computed from raw FA data. Here, we removed those reactions including DHA to avoid artifacts due to supplementation. One edge (reaction) is composed of two nodes (substrate and product), whose colors are defined as the fold changes of their abundance. Edge types including increase, decrease, and no change reflect the changes of the two associated nodes. Bars are outlined yellow if the edge type is same as the perturbation of reaction. **c**,**d**, Comparison of DHA abundance in membrane glycerophospholipids (GPLs) and diacylglycerols (DAGs) for 13 samples (**c**) before and (**d**) after substructure transformation. **e**, Proportion of samples containing a FA or substructure in 13 samples, and the associated probability distribution. **f**, The coverage in FA biosynthetic network for 13 lipid profiles using raw FAs or FA substructures. The improvement index in (**e**) and (**f**) is calculated based on the average sample or network coverage from two kinds of data. w3, omega-3; w6, omega-6; w7, omega-7; w9, omega-9; endo, endogenous; exo, exogenous; FADS1/2, fatty acid desaturase 1/2; ELOVL2/3/5/7, ELOVL Fatty Acid Elongase 2/3/5/7.

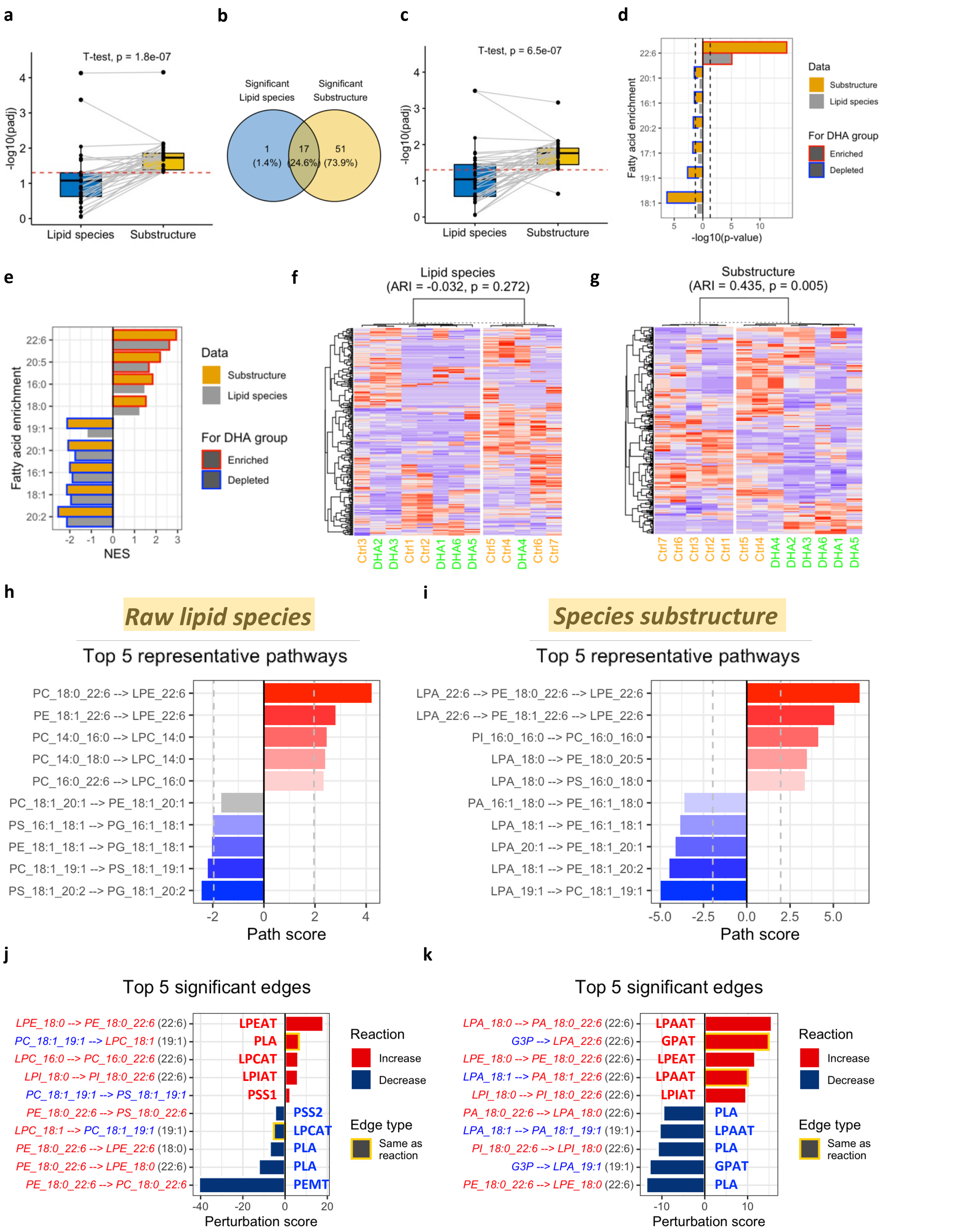


**Extended Data Fig. 2** Raw lipid species and substructure analysis for the DHA-treated dataset. **a**, Boxplots show a substantial increase in the statistical power when using the substructure method. We only include the substructures that are significant in either analysis. After missing value imputation, **b**, the Venn diagram and **c**, the boxplots also reveal a higher statistical power in the moderated t-test with the substructure method (adjusted p-value<0.05). We only include the features (lipids and substructures) that are significant in either analysis. with iLipidome (adjusted p-value<0.05) in DHA-treated RBL cells. An enrichment analysis using **d,** over representation analysis (ORA) and **e,** gene set enrichment analysis (GSEA) was further performed for significant lipid species or substructures based on FA categories. The significant enriched or depleted FA features were highlighted with a red or blue frame, respectively. **f**,**g**, Substructure profiles enable more accurate clustering on biological samples and lipid characteristics. Heatmap of hierarchical clustering for (**b**) raw lipid species and (**c**) species substructures in 13 samples. **h**,**i**, Top 5 significantly activated and suppressed representative pathways based on (**d**) raw lipid species or (**e**) species substructures. **j**,**k**, Top 5 significantly up-regulated and down-regulated edges (reactions) in the network built from (**d**) or (**e**). NES, normalized enrichment score; ARI, adjusted Rand index; GPAT, glycerophosphate acyltransferase; LPAAT, lysophosphatidic acid acyltransferase; LPCAT, lysophosphatidylcholine acyltransferase; LPEAT, lysophosphatidylethanolamine acyltransferase; LPIAT, lysophosphatidylinositol acyltransferase; PLA, phospholipase A; PEMT, phosphatidylethanolamine N-methyltransferase; PSS1, phosphatidylserine synthase 1; PSS2, phosphatidylserine synthase 2.

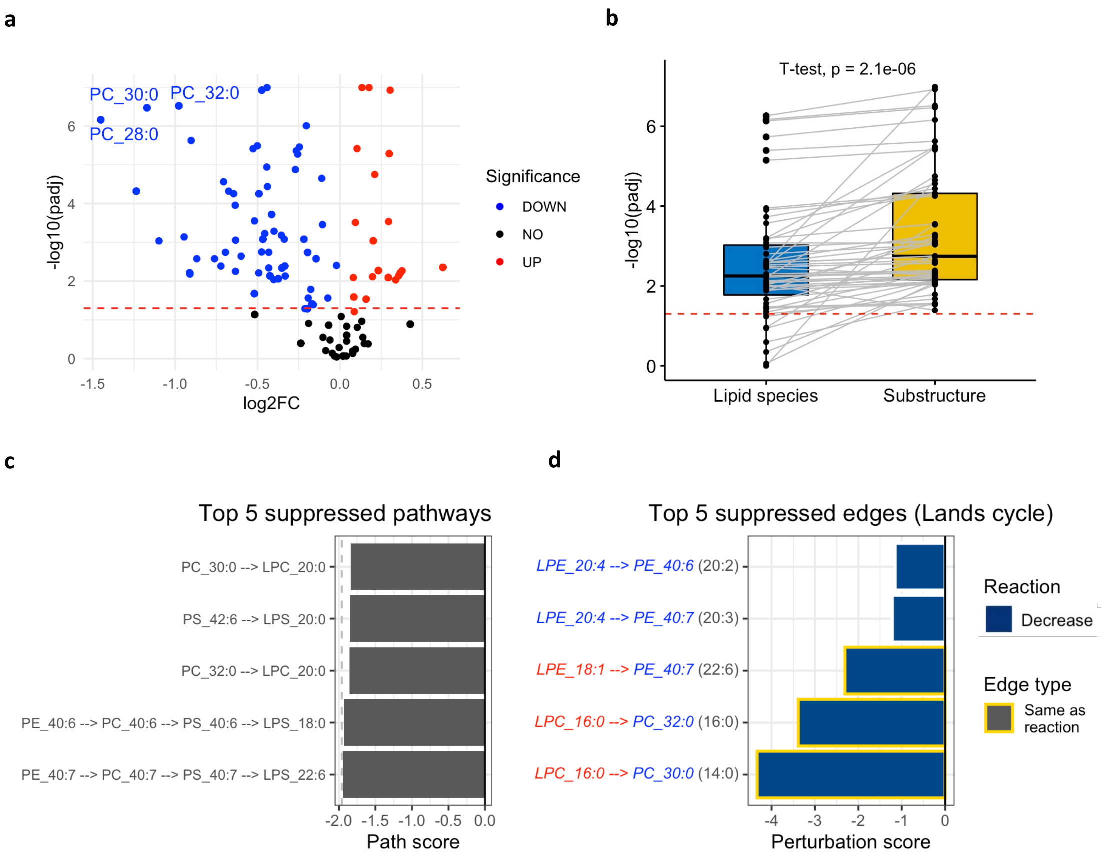


**Extended Data Fig. 3** Raw lipid species and substructure analysis for the LPCAT1 knockout dataset. **a**, The volcano plot showing the differentially expressed lipid species of U87EGFRvIII cells expressing control and LPCAT1 shRNA. n = 5 biological replicates. Two-tailed Student's t-tests with Benjamini–Hochberg correction method were used to calculate the p-values. **b**, Boxplots reveal an increased statistical power when using substructures. We only include the substructures that are significant in either analysis. **c**, Top 5 suppressed representative pathways computed from raw lipid species. **d**, Top 5 significantly suppressed edges (reactions) in the Lands cycle in the network built from (**c**).

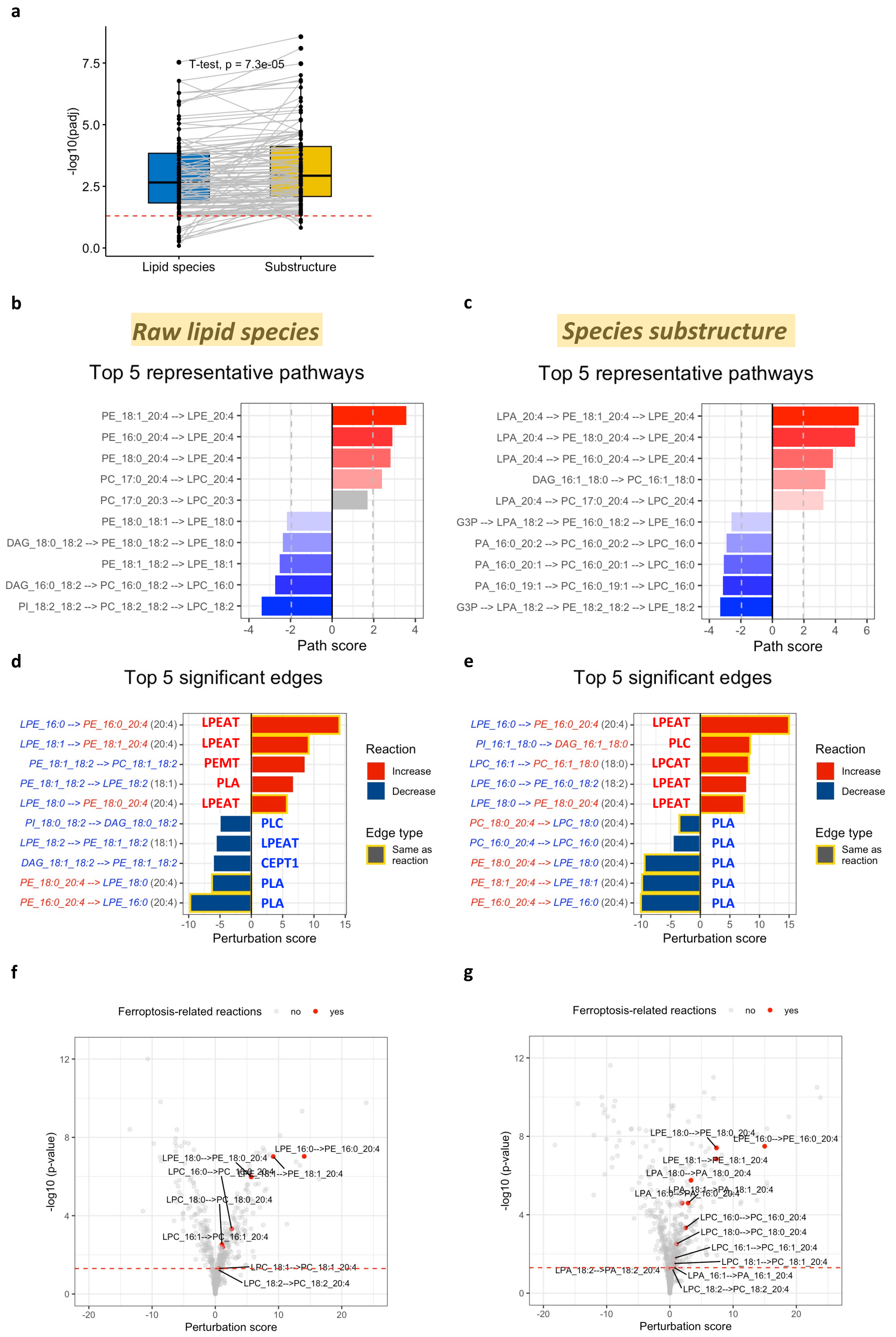


**Extended Data Fig. 4** Raw lipid species and substructure analysis for the CVD dataset. **a**, Boxplots show the improvement of statistical power in the substructure analysis. We only include the substructures that are significant in either analysis. **b**,**c**, Top 5 activated and suppressed representative pathways based on (**b**) raw lipid species or (**c**) species substructures. **d**,**e**, Top 5 significantly up-regulated and down-regulated edges (reactions) in the network built from (**b**) or (**c**). **f**,**g**, The volcano plots show all ferroptosis related reactions, such as the Lands cycle and LPAAT reactions involving arachidonic acid (AA 20:4) addition. LPCAT, lysophosphatidylcholine acyltransferase; LPEAT, lysophosphatidylethanolamine acyltransferase; PLA, phospholipase A; PEMT, phosphatidylethanolamine N-methyltransferase; PLC, Phospholipase C; CEPT, choline/ethanolamine phosphotransferase 1.

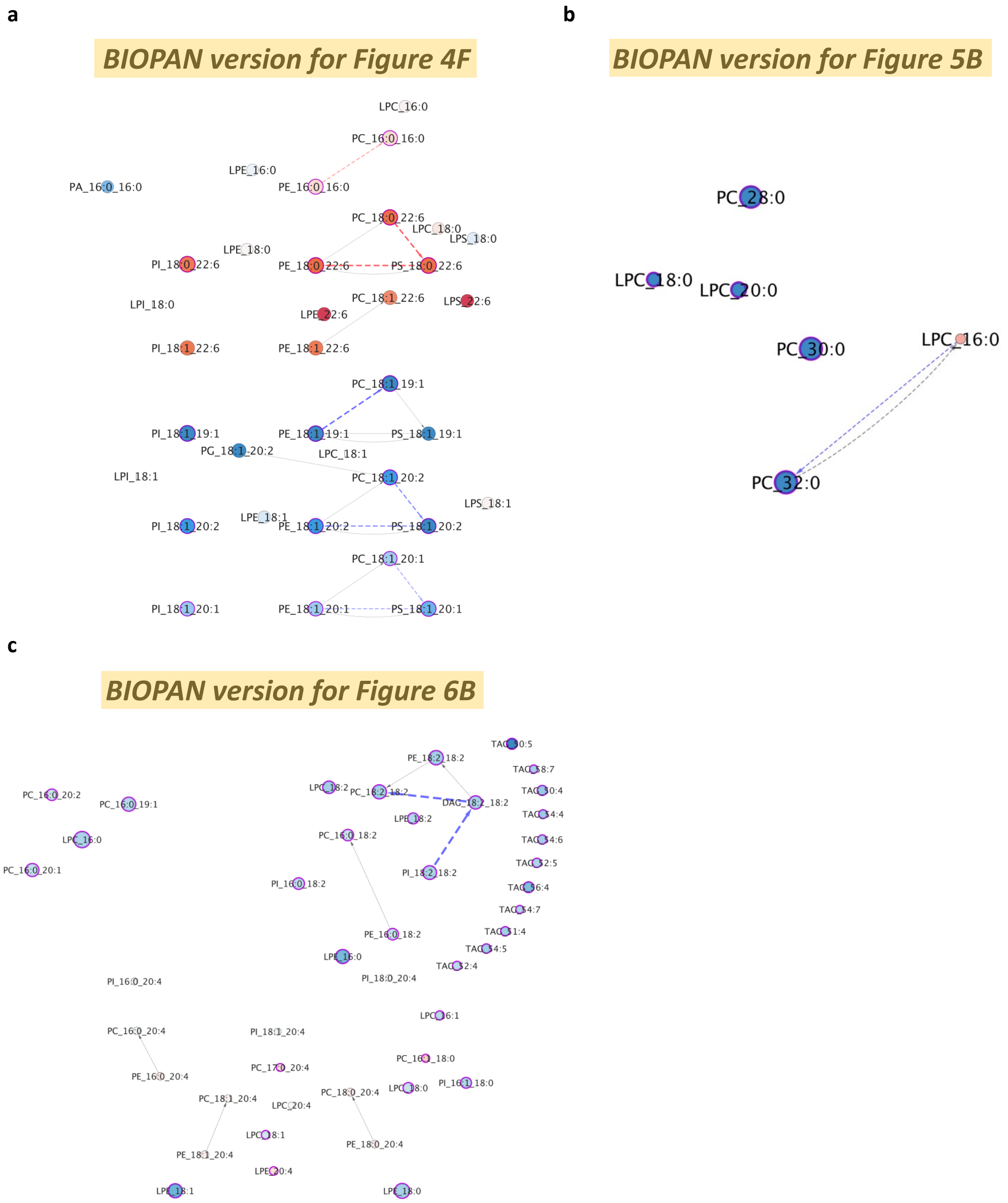


**Extended Data Fig. 5 a-c,** Application of BIOPAN's method in our dataset. The lipid species networks constructed from (**a**) the DHA-treated dataset, (**b**) the LPCAT1 knockout dataset, and (**c**) the CVD dataset based on the lipid-connecting algorithm in BIOPAN.

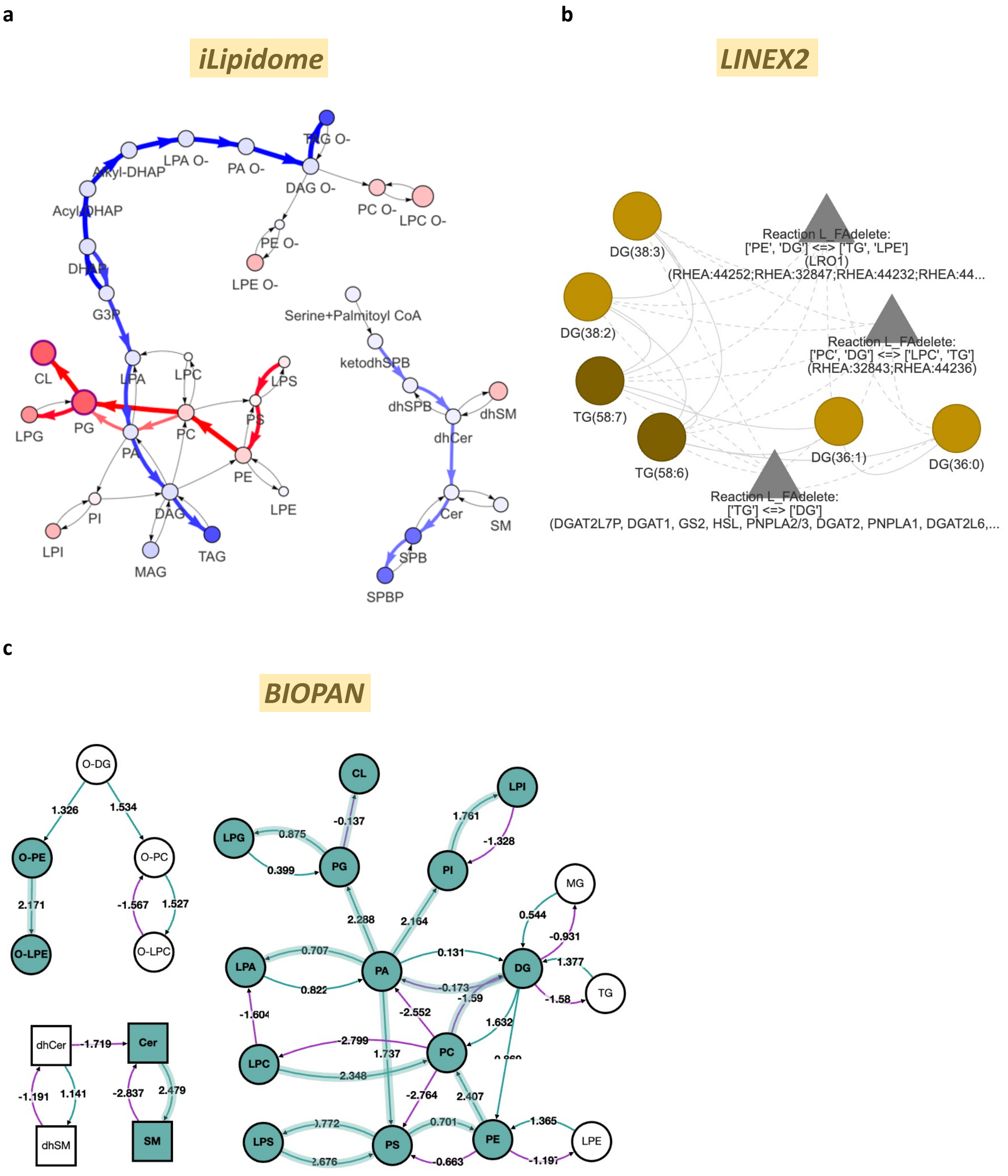


**Extended Data Fig. 6 a-c,** Comparison of the results from three tools for the dietary restriction dataset. **a**, The lipid class biosynthetic network generated in iLipidome visualizes the top 3 activated and suppressed representative pathways using red and blue colors. The intensity of the color represents the importance of these pathways. Nodes are filled according to log2 (fold change) and their sizes denote –log10 (adjusted p-value). If one node shows significant changes in abundance, its border will be marked as purple. **b**, Enriched subnetwork returned by the LINEX2's algorithm. **c**, Computed lipid class network with z-scores for active reactions in BIOPAN.

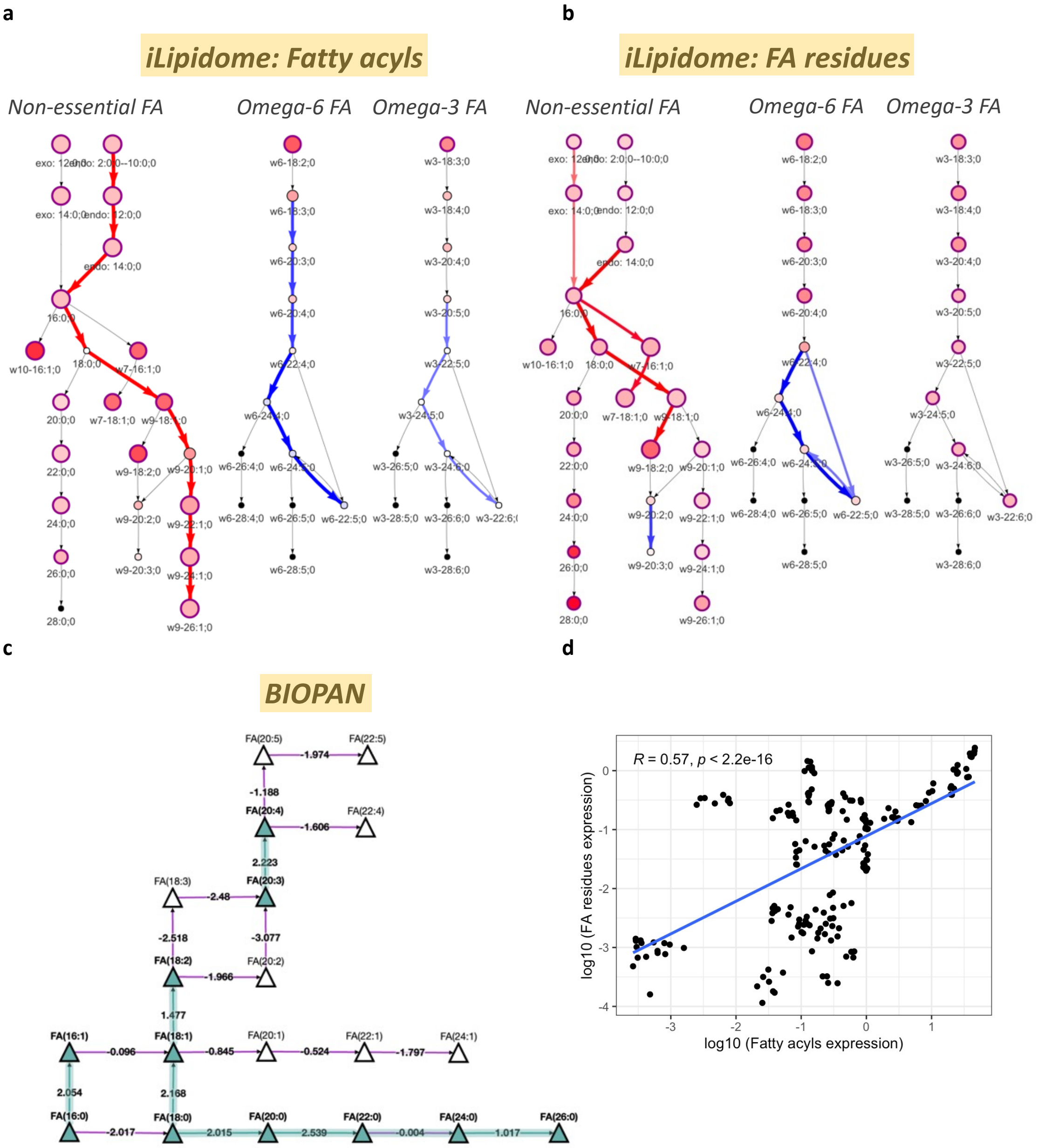


**Extended Data Fig. 7 a-d,** Comparison of the FA network from iLipidome and BIOPAN for the dietary restriction dataset. **a,b,** The FA biosynthetic network calculated by iLipidome based on the expression of **(a)** fatty acyls or **(a)** fatty acid residues extracted from other lipids. **c,** Computed FA network with z-scores for active reactions in BIOPAN. **d**, The correlation between the expression of fatty acyls and fatty acid residues.

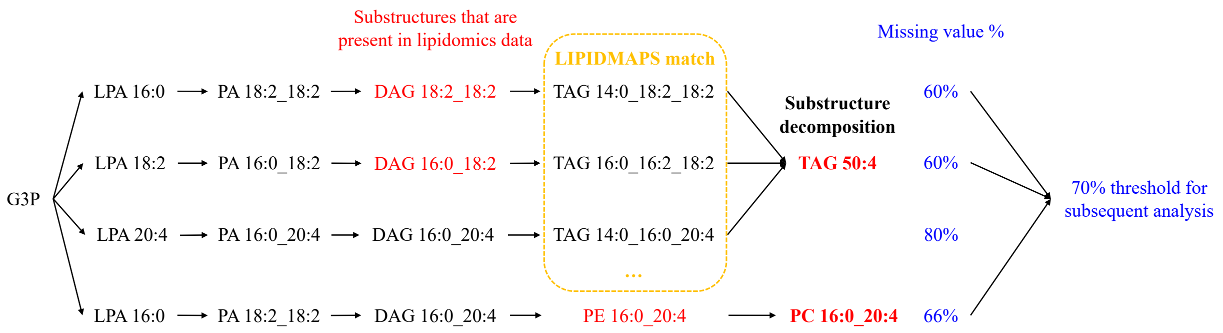


**Extended Data Fig. 8,** A workflow demonstrating how iLipidome manages substructure decomposition for the target lipids. The target lipids are represented in bold. Substructures that are present in the lipidomics data are colored in red, while missing substructures are shown in black. If the percentage of missing substructures in a particular biosynthetic pathway exceeds the specified threshold, the target lipids will not be decomposed based on this pathway (G3P→LPA 20:4→PA 16:0-20:4→DAG 16:0-20:4→TAG 50:4).

### Extended data tables

| Gene | Knockout method | Lipidomic changes found in the paper | iLipidome network type | Consistent lipidomic changes in top 5 pathways | Knockout gene in top 5 suppressed reactions |
|---|---|---|---|---|---|
| **CERS2** | Deletion | C40-44 SL↓; C34 SL↑ | SL species network | Yes | Yes (top 1) |
| **FADS3** | Deletion | SL with 0 and 2 double bonds↓; SL with 1 double bond↑ | SL species network | Yes | N/A |
| **SGMS1** | Gene trap | SM↓; HexCer↑ | SL species network | Yes | Yes (top 1) |
| **MPDU1** | Gene trap | SM↓; HexCer↑ | SL species network | Yes (dhSM) | N/A |
| **ORMDL2** | Gene trap | SM↓ | SL species network | Yes (dhSM) | N/A |
| **GNPAT** | Deletion | Ether lipids↓; PE↑ | GPL class network | Yes | No |
| **CEPT1** | Gene trap | PC, PE↓; PC O-, PE O-↑ | GPL class network | Yes | Yes (top 1) |
| **ACOT7** | Gene trap | FAs with 20 or more carbon atoms↑ | FA network | Yes | N/A |
| **DECR2** | Gene trap | Long-chain PUFAs↑ | FA network | Yes | N/A |
| **ELOVL5** | Gene trap | FAs with 18 or more carbon atoms↓ | FA network | Yes | Yes (top 1) |
| **HSD17B12** | Gene trap | FAs with 18 or more atoms↓ | FA network | Yes | Yes (top 1) |

**Extended Data Table. 1,** The overall performance of iLipidome in identifying the essential pathways and reasons using a multiple gene knockout dataset. N/A means the knockout genes cannot be found in our reference lipid biosynthetic pathways. CERS2, Ceramide Synthase 2; FADS3, Fatty Acid Desaturase 3; SGMS1, Sphingomyelin Synthase; MPDU1, Mannose-P-dolichol Utilization Defect 1 Protein; ORMDL2, ORMDL Sphingolipid Biosynthesis Regulator 2; GNPAT, Glyceronephosphate O-Acyltransferase; CEPT1, Choline/Ethanolamine Phosphotransferase 1; ACOT7, Acyl-CoA Thioesterase 7; DECR2, 2,4-Dienoyl-CoA Reductase 2; ELOVL5, ELOVL Fatty Acid Elongase 5; HSD17B12, Hydroxysteroid 17-Beta Dehydrogenase 12; SL, sphingolipod; SM, sphingomyelin, HexCer, hexosylceramide; PE, phosphatidylethanolamine; PC, phosphatidylcholine; PE O-, ether-linked phosphatidylethanolamine; PC O-, ether-linked phosphatidylcholine; FAs, fatty acids; PUFAs, polyunsaturated fatty acids; GPL, glycerophospholipid; dhSM, dihydrosphongomyelin.

| | iLipidome | BIOPAN | LINEX2 |
|---|---|---|---|
| **Tool type** | R function | Web tool | Web tool |
| **Display result** | Active/Suppressed pathway and reaction (gene) | Active/Suppressed pathway | Enriched lipid subnetwork |
| **Calculation basis** | Substructure-based statistics | Ratio of product over substrate | Substrate-product change |
| **Parse FA for lipids** | Automatic mapping to LIPIDMAPS database | Require FA data | Require FA data |
| **Identify lipid intermediate/precursor** | Yes | No | No |
| **Improve statistical power and data sparsity** | Yes | No | No |
| **Network extension** | Yes | No | Yes |

**Extended Data Table 2,** Comparison of tools for the network analysis of lipidomics data.
